# A method for efficient, rapid, and minimally invasive implantation of individual non-functional motes with penetrating subcellular-diameter carbon fiber electrodes into rat cortex

**DOI:** 10.1101/2025.02.05.636655

**Authors:** Joseph G. Letner, Jordan L. W. Lam, Miranda G. Copenhaver, Michael Barrow, Paras R. Patel, Julianna M. Richie, Jungho Lee, Hun-Seok Kim, Dawen Cai, James D. Weiland, Jamie Phillips, David Blaauw, Cynthia A. Chestek

**Author notes:** **Declaration of Interests** JDW has a financial interest in PtIr material. All other authors have no conflicts of interest to disclose.

## Abstract

**Objective:** Distributed arrays of wireless neural interfacing chips with 1-2 channels each, known as “neural dust”, could enhance brain machine interfaces (BMIs) by removing the wired connection through the scalp and increasing biocompatibility with their submillimeter size. Although several approaches for neural dust have emerged, a procedure for implanting them in batches that builds upon the safety and performance of currently used electrodes remains to be demonstrated.

**Approach:** Here, we demonstrate the feasibility of implanting batches of wireless motes that rest on the cortical surface with carbon fiber electrodes of subcellular diameter (6.8-8.4 µm) that penetrate to a target brain depth of 1 mm without insertion aids. To simulate their implantation, we assembled more than 230 mechanically equivalent motes and affixed them to insertion tools with polyethylene glycol (PEG), a quickly dissolvable and biocompatible material. Then, we implanted mote grids of multiple configurations into rat cortex *in vivo* and evaluated insertion success and their arrangement on the brain surface using photos and videos captured during their implantation.

**Main Results:** When placing motes onto the insertion device, we found that they aggregated in molten PEG such that the array pitch was only 5% wider than the dimensions of the mote bases themselves (240 x 240 µm). Overall, we found that motes with this arrangement could be inserted into rat cortex with a high success rate, as 171/186 (92%) motes in 4x4 (N=4) and 5x5 (N=5) square grid configurations were successfully inserted using the insertion device alone. After implantation, measurements of how much motes tilted (22±9°, X̄±S) and had been displaced relative to their original positions were smaller than those measured for devices implanted inside the brain in the literature.

**Significance:** Collectively, these data establish the viability of safely implementing motes with ultrasmall electrodes and epicortically-situated chips for use in future BMIs.

## Introduction

Neural interfaces for recording and stimulating the nervous system have been used to study and treat a wide range of conditions [1]. These include brain machine interfaces (BMIs), which have been used to restore functions lost by neurological injury or disease [2,3]. Successful applications of BMIs include control of prosthetic arms [4,5], control of a computer [6,7], speech restoration [8,9], vision restoration [10,11], walking [12], paralyzed limb reanimation [13], and the list is expected to grow. Despite successful multi-year clinical trials [3] and the recent regulatory success of newer systems, such as those made by Synchron, Neuralink, and Precision Neuroscience [14], BMIs have yet to gain the approval required to become a broadly applicable clinical treatment [15]. To become a viable clinical solution, challenges associated with, but not limited to, BMI hardware must be mitigated [16]. The most pressing of these challenges is expanding the number of electrode channels in a safe and manageable way so that BMIs can interact with more neurons [17].

Large networks of neural interfacing chips that are both submillimetric and wireless could potentially solve the hardware issues persisting in BMIs [18]. Often called “neural dust” after one type of these devices [2,19], these chips are comprised of a single electrode and an application specific integrated circuit (ASIC) responsible for wirelessly harvesting power and communicating while performing intended functions, such as stimulating or digitizing recorded neural signals. The wireless link removes a wired connection that is currently transcranial and commonly transcutaneous. This could be advantageous for the BMI user because the risks of infection [20] and foreign body responses (FBRs) from micromotion [21] could be reduced. This wireless link would also obviate the physical tether to a computer system, offering the user more freedom of movement [22]. Additionally, restoring naturalistic hand control through BMIs may necessitate recording from as many as hundreds of thousands of neurons [23]. At such channel counts, wired connectors become increasingly difficult to manage [24,25]. Even with 100 channels, the wired connection to the array is already a frequent source of device failure [26,27]. Without a wired connection, neural dust electrodes are effectively individualized and can be implanted in modular batches to increase coverage or more efficiently target brain regions of interest [21,24]. Also, if they are sufficiently small, they may cause less damage and be more biocompatible than current arrays [28,29]. Collectively, these potential benefits have garnered interest in this architecture and a large increase in innovative approaches [18,19,30–34].

BMIs often target neurons deep in cortex to obtain high-resolution neural firing patterns [2,35]. However, implanting neural dust to these cortical depths in a way that improves upon the safety and performance of existing electrodes is difficult due to the size of the ASICs in existing designs. Placing neural dust onto the brain surface will primarily record field potentials rather than spikes from individual neurons [36]. Some groups have proposed implanting the entire device into the brain [31,37,38]. However, these devices have dimensions comparable to traditional silicon probes [39,40], which are widely known to induce a negative FBR and damage surrounding cells over time [41–43]. As a result, histology of implant sites associated with inserting neural dust into cortex has pointed to a similar FBR [37,44]. It would be better to place the bulk of the device onto the brain surface instead, and use a smaller electrode extending from the chip to penetrate the brain and interface with neurons directly.

In order to safely implant neural dust with this geometry, the electrodes will need to be sufficiently small and biocompatible to avoid eliciting the FBR observed with traditional penetrating electrodes. Prior work using epicortical dust with large wire electrodes (≥50 µm diameter) to reach the target depth [33,45–47] were not ideal as microwire electrodes of this size can also elicit a negative FBR [48,49]. Instead, using penetrating electrodes that have reduced bending stiffness and electrode dimensions that are cellular scale or smaller can minimize the damage to surrounding tissue [28,50]. Many ultrasmall probes have recently exploited these principles to achieve a markedly reduced FBR [51]. These include the nanoelectronic thread (NET) electrodes [52,53], flexible polymer electrodes [54], mesh electrodes [55,56], and carbon fibers [57,58]. However, the higher biocompatibility of ultrasmall electrodes typically comes at the cost of increased difficulty when inserting them into the brain, requiring one or more insertional aids [59]. Implanting a network of thousands of individual electrodes further compounds this difficulty [27] and highlights the need for a biocompatible penetrating electrode that is easy to insert.

Of the highly biocompatible penetrating electrodes listed above, subcellular-scale carbon fiber electrodes (6.8 µm) may be the easiest to insert into the brain, due mainly to carbon’s strength at small size [60]. Our group has previously shown that carbon fibers with chemically sharpened tips reliably insert at least 1.5 mm into the brain without any insertional aid [61]. We have also proposed [62] a neural interfacing system comprised of hundreds of wireless motes, each with its chip situated on the brain surface and a carbon fiber electrode penetrating to the targeted cortical depth (Fig. 1). These motes would harvest power from near-infrared (NIR) light emitted by a repeater embedded in the skull and relay messages back to the repeater via light emitting diodes (LEDs) blinking packets of data [63,64]. This repeater would, in turn, communicate with a receiver externally placed on the scalp for interfacing with a computer (Fig. 1a). Currently, this proposed system has two versions: a mote intended for recording neural signals and another for stimulating neurons, called the ReMote and StiMote, respectively (Fig. 1b.). Common to both designs is the integration of two constituent chips: one composed of a photovoltaic cell and LED for wirelessly powering and communicating, and one housing an ASIC responsible for the mote’s function. Many of the system’s components have been successfully validated with prototypical components [40], including the ReMote ASIC [39], the StiMote ASIC [40], optical uplink [63], power harvesting [65], RF wireless uplink [66], and carbon fiber electrode arrays [60,61,67]. A critical aspect of the translation of the mote approach is a paradigm for inserting arrays of motes in batches. Therefore, in this work, we sought to assess methods for the insertion of large quantities of motes by using non-functional analogs of dimensions similar to the StiMote (Fig. 1c).

**Figure 1.**
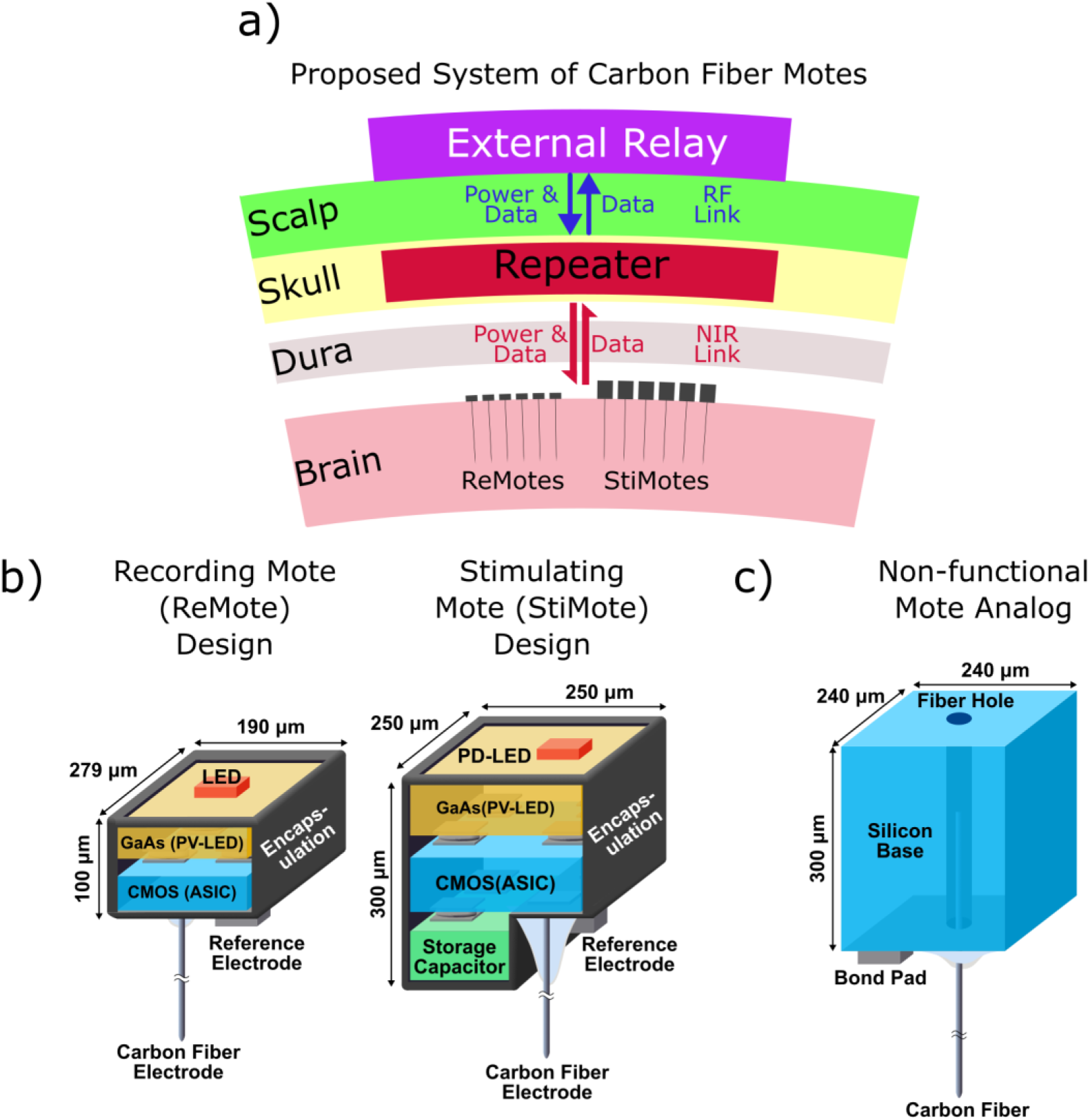
Carbon fiber mote system and proposed mote designs. a) Conceptual diagram showing a network of carbon fiber motes functioning in brain. A network of ReMotes and/or StiMotes for recording or stimulating neurons, respectively, with chips sitting on the brain are powered by and communicate via a repeater unit embedded in the skull via near infrared (NIR) light. Carbon fibers penetrating the brain reach the targeted cells. The repeater is powered by and communicates via radiofrequency (RF) with an interface externally placed on the scalp. b) Cross-sectional diagrams of the proposed ReMote (left) and StiMote (right). For both designs, the top chip has a photovoltaic (PV) cell and light-emitting diode (LED) for wireless functionality, and an application-specific integrated circuit (ASIC) chip for device functionality below the PV-LED chip. The StiMote also has a capacitor for charge storage. Both devices have a carbon fiber electrode protruding from the bottom face and a conductive pad for referencing. c) Non-functional mote analog used in this study, designed to have similar dimensions to proposed mote designs. A more detailed layer-by-layer diagram for the analogs can be seen in Figure S1. Adapted from [40].

Here, we show the fabrication of carbon fiber motes to determine viable carbon fiber assembly and implantation techniques for future wireless designs. In this work, we produced more than 230 non-functional mote analogs consisting of a micromachined silicon base with an extruding carbon fiber. These devices were used to introduce and test a method for swiftly, efficiently, easily, and simultaneously implanting up to 25 motes in rats with epicortically-situated chips and minimally-invasive carbon fiber electrodes that penetrate 1 mm in cortex. Implanting motes using this scheme into rat cortex *in vivo* (N=9 implantations) showed a 92% success rate (171 of 186). Additionally, implanted motes had lower changes in spatial arrangement (22±9° tilt angle, 65±55 µm displacement) compared to intracortically implanted designs as reported in the literature. To our knowledge, this work introduces the first successful method for implanting epicortically situated neural dust in batches and provides the first step in translation to clinical use.

## Methods

### Fabrication of silicon bases for non-functional carbon fiber motes

Silicon bases were fabricated using standard cleanroom micromachining processes. These bases were designed to be prismatic with dimensions (240 x 240 x 300 µm, L x W x H) similar to proposed functional carbon fiber motes (Fig. 1b). The hole through the base, used for holding a protruding carbon fiber electrode, was designed to be conductive with connection to a backside metal pad for use as a wired mote. Ultimately, we did not add wires for experiments that would require conductive devices. All steps are listed here anyway to provide a full account of the motes’ material composition. Figure S1 shows a cross-sectional diagram of the non-functional mote and comprising layers.

The source substrate for silicon bases was a 4” p-doped silicon wafer that was 300 µm thick and polished on both sides (2345, UniversityWafer, South Boston, MA). The wafer was coated in a layer of Parylene C approximately 5 µm thick (PDS2035CR, Specialty Coatings Systems, Indianapolis, IN) with AP-174 adhesion promoter. This layer was applied to the wafer backside to later hold the individuated mote analogs. The layer of Parylene C on the wafer’s topside was removed via oxygen plasma ashing (YES-CV200RFS(E), Yield Engineering Systems, Fremont, CA). SPR 220 (3.0) Photoresist (MEGAPOSIT SPR 220 (3.0), Series i-Line; Dow Chemical Co., Midland, MI) applied to the topside followed by deep reactive ion etching (DRIE) (STS Pegasus 4, SPTS Technologies Inc., Newport, United Kingdom) produced a 20, 25 or 30 µm hole in each device. These three hole sizes intentionally differed for determining viable carbon fiber hole sizes. Due to inconsistencies in the DRIE, the hole did not always go through the whole device. A 30 nm passivation layer of Al_2_O_3_ followed by a 10 nm conductive layer of TiN were added to the topside and the hole sidewalls via atomic layer deposition (Fiji ALD, Veeco Instruments Inc., Plainview, NY) to render the hole and topside electrically connected. A 50 x 50 µm contact pad (10 nm Ti, 200 nm Au) was sputtered (Lab 18, Kurt J. Lesker Co., Pittsburgh, PA) onto the topside beside the hole and patterned via the liftoff process, patterned with LOR 10B (Kayaku Advanced Materials Inc., Westborough, MA) followed by SPR 220 (3.0). Singulating the sample into individual mote bases, similar to wafer dicing, was then achieved in three steps: 1) patterning with SPR 220 (3.0) photoresist, 2) portions of the TiN and Al_2_O_3_ layers that covered intended etch lanes for the silicon bases were etched via ion milling (Nanoquest II, IntIvac Thin Film Corp., Fort Collins, CO) 3) the sample was mounted to a 4” carrier wafer with Santovac 5 (Santolubes, LLC; Spartanburg, SC) and singulated via DRIE (STS Pegasus 4). Devices were held in place on the layer of Parylene C for storage indefinitely until removed for assembling a non-functional carbon fiber mote analog (Fig. 3a).

### Assembly of non-functional carbon fiber mote analogs

A process flow diagram of the assembly steps is shown in Figure S2.

#### Affixing carbon fibers to mote bases

Non-functional motes were first assembled in batches that started with 15-20 motes. Mote bases were pulled off of the source wafer with forceps (Fig. 3a). In most cases, Parylene C remained on the bottom side of the base, which was scraped off with a razor blade (11-515, Stanley Hand Tools, New Britain, CT). Next, 10,000 MW polyethylene glycol (PEG) (309028, Sigma Aldrich, St. Louis, MO) was placed onto an unused glass coverslip (12-544-E, Fisher Scientific, Waltham, MA), then melted and spread via a wire wrapped around a soldering iron set to 350-400° F (∼177-204° C) [60] along the middle of the coverslip (Fig. 3b). Mote bases were placed onto the PEG layer with the topside facing up. Once again using the soldering iron wrapped in wire, the surrounding PEG was melted so that the base of the motes would sink and become flat against the coverslip. This positioning ensured that the topside was level during fiber placement. Mote bases were placed with enough spacing between them to prevent forceps from hitting fibers in neighboring motes. Epoxy (Epo-Tek 301, Epoxy Technology, Billerica, MA) was deposited onto the topside via a pulled glass capillary. A 4-6 mm carbon fiber (T-650/35 3 K, Cytec Thornel, Woodland Park, NJ) was threaded into the base’s hole. Successful placement of the fiber generally required inserting at an angle and bending the fiber slightly (approx. 30°) so that when the fiber straightened after releasing tension, it would be pushed deeper into the hole (Fig. S3). The epoxy was partially cured overnight at room temperature and then oven-cured at 140°C for 20 minutes [68]. In later batches, the bonds between the base and fiber were evaluated for stability by bending or pulling the fibers with forceps. An additional layer of 301 epoxy was added to devices with unsecure fibers and oven-cured shortly afterward (140°C, 20 minutes). Motes that received additional epoxy and still failed to become sufficiently secure were not used.

#### Encapsulation in Parylene C

A subset of motes (N=105) was encapsulated in Parylene C [47,60]. First, motes were released from the securing PEG by either holding them with forceps under heat or by placing the entire coverslip into a petri dish filled with deionized (DI) water and waiting for the PEG to dissolve. In parallel, silver epoxy (H20E or H20S, Epoxy Technology) was placed onto a printed circuit board (PCB) [69] and shaped to have a dune-like slope [68] to be able to hold the motes by their carbon fibers (Fig. 3c). The epoxy was oven-cured at 140°C for at least 20 minutes [60], after which 660–760nm of Parylene C was deposited (PDS2035CR) [67]. A glass slide was included during deposition to be able to measure the Parylene C thickness, which was performed using a stylus profilometer (DektakXT, Bruker Corp., Billerica, MA).

#### Fire-sharpening the tips of carbon fiber motes

Prior to tip sharpening, the fibers were cut to a length of 1050 µm with microsurgical scissors (15003-08 or 15002-08, Fine Science Tools, Foster City, CA) [60], where the extra 50 µm acted as a buffer that would be burned away during sharpening. During cutting, the coverslip or glass slide holding the devices was placed on its side, held by putty alone or onto the side of a wooden block via putty and the fiber length was approximated with an eyepiece reticle in the stereoscope. Generally, motes with fibers of lengths ±50 µm of the target length were kept.

To enable fibers to penetrate the brain without insertion aids, tips were fire-sharpened using a modified procedure reported by Welle *et al.* [70] (Fig. 3d). First, a strip of parafilm (PM996, Bemis Company, Neenah, WI) was cut and placed onto a glass slide (12-550-15, Fisher Scientific) being heated by a PCB warmer (CSI358, Circuit Specialists, Tempe, AZ). Once the parafilm turned clearer from heating, a gloved finger was run across it to straighten the layer, and motes were then partially stuck into the parafilm layer with fibers facing up. The slide was placed into a dish filled with DI water and the water level adjusted so that fiber tips were touching the meniscus. Levelling the water was visualized with a pen camera (TSMS100, Teslong, Shenzhen, China). A butane microtorch with a 1-3 cm long flame was waved along the water such that the portion of the flame that was hotter at the center briefly pushed down the meniscus of the water, exposing the fibers to the flame directly. This was repeated several times before visually inspecting the tip shape through a benchtop microscope. Due to the variable heights of the tips from deviations in fiber angle and from different depths in the parafilm layer, often a subset of fibers would sharpen first, and would then be removed to prevent oversharpening. The process of torching, inspection, and removal was repeated until all fibers in a batch were sharp.

#### Assembling insertion devices and mote grids

Insertion devices were assembled using the following steps. First, the head of a nail with a shaft diameter of 1.4 mm was cut off with wire cutters. The cross-section produced by the cut was filed or sanded so that it was approximately orthogonal to the shaft’s length. A glass coverslip was cleaved into a square shape using either a dressing stone or tungsten carbide scribe pen and a stripped wire. For 3x3 and 4x4 mote analog grids, the target size was 1.5 x 1.5 mm, while for 5x5 grids, the intended size was 2 x 2 mm. A vial clamp then held the nail in place with the cross-section pointing upward. Initially, a small layer of Norland Optical Adhesive 61 (NOA 61, Norland Products, Inc., Cranbury, NJ) was applied to the cross-section, the coverslip fragment placed and pressed onto the epoxy, and the epoxy cured using ultraviolet light. In later iterations, Epo-Tek 301 epoxy was used instead, and heat cured for 20 minutes at 140°C [68].

Mote grids were assembled by placing the motes onto the glass portion of the insertion device and held with PEG. Figure 2a shows a conceptual diagram of a completed grid. During mote placement, the insertion nail was held upright with a vial clamp such that the glass was on top and also was parallel to the floor. Small flakes of PEG were placed onto glass and melted by firmly touching the nail under the glass with a soldering iron set to 350-400° F (∼177-204° C). One or two non-functional motes were carried by forceps onto the insertion device near its edge. With forceps in one hand and the soldering iron in the other, the iron was then pressed against the nail to melt the PEG while the forceps ensured that fibers were orthogonal to the glass surface. If sufficient PEG was present, motes would be pulled toward other motes already present due to surface tension. These steps were repeated until the desired number of motes was present. Periodically, motes were guided in the melted PEG with forceps to facilitate the desired arrangement (e.g., 4x4 square grid). Completed grids on the insertion device were stored horizontally, and could be stored for long periods of time (one unused grid had been stored for at least 16 months when drafting this manuscript). After implantation, if the insertion nail was still viable, it was saved for reuse.

**Figure 2.**
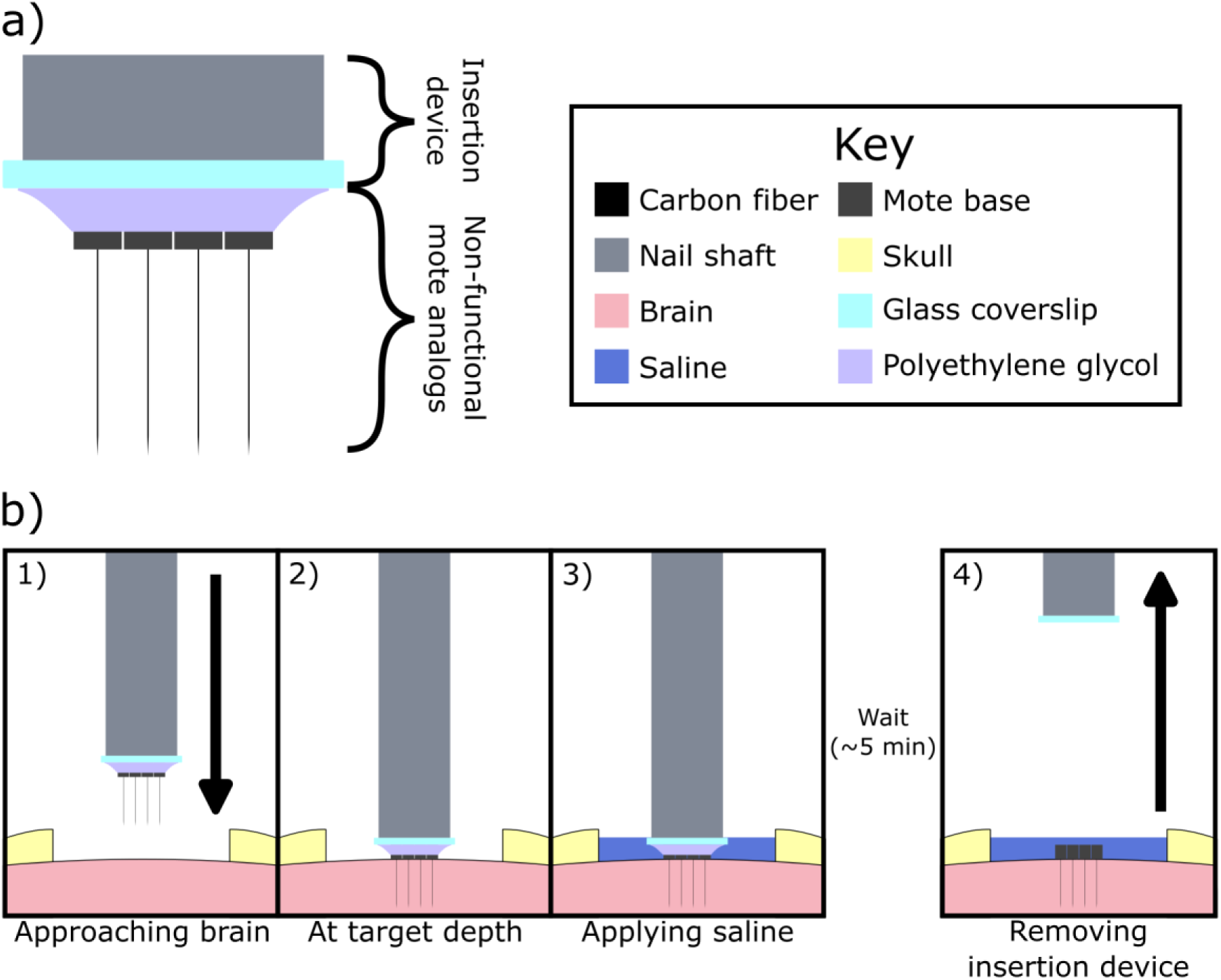
Diagram of carbon fiber mote insertion. a) Cartoon profile of a completed mote grid ready for implantation, with motes held to the device by dissolvable polyethylene glycol (PEG). b) Procedure for mote implantation shown in four panels: 1) The devices are lowered to the brain. 2) Driving the insertion device downward ceases once the fibers have penetrated the brain and reached the target depth, which is indicated by the bases having reached the brain’s surface. 3) Sterile saline is applied to the craniectomy to dissolve the PEG. 4) After waiting for a short time (∼5-7 minutes), the insertion device is removed from the craniectomy, leaving the motes in place.

In N=1 4x4 mote grid, N=10 motes were reused after a previous implantation (these were from insertion 8, reused in insertion 11). Prior to placing these motes in a mote grid, they were soaked overnight in an enzymatic detergent solution (approximately 1.5% in deionized water, vol/vol) (Enzol, 2252, Advanced Sterilization Products, Irvine, CA). Also, a 4x4 grid that had been sterilized with ethylene oxide gas was upgraded to a 5x5 with new motes (insertion 14). For N=2 mote grids, additional PEG was added at the mote bases to increase stability during implantation after observing that premature PEG release was a failure mode.

#### Preliminary mote analog devices

Many of the assembly methods listed above were developed while assembling the first batches of non-functional motes (N=45 motes) used in proof-of-concept insertions (Fig. S4, Table S1). These preliminary devices are largely excluded from the main text and relegated to the supplementary information, with the exception of reporting their insertion success rate, and including their failure modes in Table 2 and its associated text. The steps were mostly the same, but none of these preliminary motes were coated in Parylene C nor were the tips fire-sharpened. Because the tips were blunt, the fibers for preliminary devices were cut to a target length of 500 µm. Some of these devices were also held in place using 2050 MW PEG (295906, Sigma Aldrich) instead, a higher temperature setting on the soldering iron was used, and the fibers were dipped in DI water prior to insertion to remove excess PEG. In one batch, 301 epoxy and 353ND-T (Epoxy Technologies) were incorrectly mixed together, but fibers were sufficiently fastened for insertion testing.

### Implantation of non-functional carbon fiber mote analogs

The mote implantation procedure was performed in N=12 male rats weighing 341-628 g where N=5 rats were Sprague-Dawley and N=7 rats were Long-Evans strains. The surgical steps for both non-survival (acute) and survival (chronic implant) procedures followed previously published methods [69,71] up to the steps specific to mote implantation. If performing a survival surgery, mote grids were sterilized via ethylene oxide gas sterilization (N=5), where N=1 grid was sterilized via a vendor (Life Science Outsourcing, Inc., Brea, CA), and sterile surgical technique was followed. General anesthesia was provided with isoflurane at 5% (v/v) and 1-3% (v/v) for maintenance. Carprofen (5 mg/kg) was injected subcutaneously as an analgesic. During the procedure, temperature, breath rate, and anesthetic depth were continually monitored. The animal was transferred to a rat stereotaxic frame to ensure head stability throughout the procedure. The scalp was shaved and sterilized with 1% betadine and then 70% ethanol (v/v) before performing a midline incision starting between the eyes and ending at the posterior end of the skull. Tissue covering the skull was pulled away with hemostats, and the skull cleaned with 3% hydrogen peroxide, followed by saline. A skin marker pen was used to mark intended craniectomy boundaries and, if applicable, bone screw coordinates. If the procedure was intended for chronic implantation, a surgical drill was used to drill 6-7 tap holes for screwing in stainless steel (19010-00, Fine Science Tools, N=3 rats) or polyetheretherketone (PEEK) (PKMCK164, High Performance Polymer, Didcot, United Kingdom, N=1 rats) bone screws. Bone screws (1ZY93, Grainger, Lake Forest, IL) were also implanted during N=1 non-survival implant. A craniectomy was then made with a surgical drill and rongeurs. Approximate stereotaxic locations of each craniectomy along with its shape are listed in two tables. The first five insertions (N=3 rats) were performed with preliminary mote devices (3x3 grids, 500 µm long fibers with blunt tips). Craniectomy details for these insertions are listed in Table S1. Craniectomy details for the remaining insertions (4x4 & 5x5 grids, 1 mm long fibers with sharpened tips) are listed in Table 1. To record videos of the insertion from the side, 1-2 pen cameras (TSMS100, Teslong), held by the provided stands, vial clamps, or Flexbars (18032, Flexbar Machine Corporation, Islandia, NY), were placed facing the craniectomy and controlled by laptops. The shaft of the insertion device was then fastened to an electrode holder (M3301EH, World Precision Instruments, Sarasota, FL) held by the manipulator arm of the stereotax frame, and then lowered to the position where at least one carbon fiber touched the dura mater. After zeroing the dorsoventral axis at this position, the arm was retracted upward and the dura was removed with a bent 23G needle and/or microsurgical scissors. During steps that did not include interaction with the craniectomy, such as setting up cameras and fastening the mote grid, gel foam (SP40400, Braintree Scientific, Braintree, MA) soaked in sterile saline was placed into the craniectomy to avoid desiccation.

**Table 1.** Insertion experiment details. Mote insertion experiments used various configurations of mote grids in several areas of rat cortex. The first five insertions in N=3 rats were performed with preliminary devices, and surgical details for those experiments are listed in Table S1. Relevant experimental details are listed here for more information on how these experiments differed. Craniectomy coordinates are relative to bregma. Abbreviations: Timespan column: A=Acute, C=Chronic. Hemisphere column: L=Left, R=Right. Craniectomy coordinates column: M/L=Medial/Lateral. A/P=Anterior/Posterior. Craniectomy Shape column: T=Trapezoid, R=Rectangle, C=Circle. Evaluation & Success Rate columns: S=Success, F=Failure, E=Exception. Note: insertion 13 was excluded due to surgeon error, see the Methods section for details.

| Insertion # | Rat # | Timespan (A/C) | Layout | Reused Motes (#) | Coated in Parylene C (Y/N) | Hemisphere | Craniectomy Coordinates |  | Craniectomy Shape | Evaluation |  |  | Success Rate (%) |
| --- | --- | --- | --- | --- | --- | --- | --- | --- | --- | --- | --- | --- | --- |
|  |  |  |  |  |  |  | M/L | A/P |  | S | F | E |  |
| 6 | 4 | C | 4x4 | 0 | Y | L | 1.0-4.5 | (-7.5)-(-4.5) | T | 15 | 1 | 0 | 94 |
| 7 | 5 | C | 4x4 | 0 | Y | L | 1.0-4.5 | (-7.5)-(-4.5) | T | 16 | 0 | 0 | 100 |
| 8 | 6 | A | 5x5 | 0 | N | R | 0.5-4.0 | 2.5-6.5 | R | 22 | 0 | 3 | 100 |
| 9 | 7 | A | 5x5 | 0 | N | L | 0.5-4.0 | (-3.5)-(-0.5) | R | 25 | 0 | 0 | 100 |
| 10 | 8 | C | 4x4 | 0 | Y | R | 0.5-3.0 | 1.5-4.0 | C | 14 | 2 | 0 | 88 |
| 11 | 9 | A | 4x4 | 10 | N | R | 1.5-4.0 | (-6.0)-(-3.0) | C | 11 | 5 | 0 | 69 |
| 12 | 10 | A | 5x5 | 0 | N | R | 1.0-4.0 | 2.0-5.0 | R | 20 | 5 | 0 | 80 |
| 14 | 12 | A | 5x5 | 0 | Y | L | 1.0-4.5 | (-5.0)-(-1.0) | R | 24 | 1 | 0 | 96 |
| 15 | 12 | A | 5x5 | 0 | N | R | 0.5-4.5 | (-5.5)-(-1.5) | R | 24 | 1 | 0 | 96 |
| **Total** | - | - | - | - | - | - | - | - | - | 171 | 15 | 3 | 92 |

At this point, the steps specific to mote implantation were performed, which are shown diagrammatically in Figure 2b (see Figure 6 for surgical photos). The pen camera(s) started recording. The procedure was also recorded through the surgical scope (OPMI CS-NC, Carl Zeiss AG, Oberkochen, Germany) using an attached camera (HVD-37A, Hitachi, Tokyo, Japan: N=5 insertions; MU1803-HS, AmScope: Irvine, CA: N=10 insertions) to record the surgeon’s point of view. The stereotaxic arm was quickly lowered back toward the zero point to approach the brain (Fig. 2b, panel 1). All movements of the stereotaxic arm were controlled by manually adjusting the knobs. Once the fibers started touching the brain, the insertion speed was slowed. Sometimes, the motes were retracted a short distance to improve penetration and allow the brain surface to relax. If insertion had started and driving the fibers down revealed that there was insufficient room for the insertion device’s glass coverslip or fibers were not penetrating, the probe was repositioned and insertion restarted. The device was driven ventrally to a point where the silicon bases touched the brain, and were generally pushed down more to ensure a complete insertion (Fig. 2b, panel 2). Sterile saline was then applied via an injection needle to dissolve the PEG (Fig. 2b, panel 3) for at least 5-7 minutes. For proof-of-concept insertions, the dissolution time was not standardized and was typically shorter. In most insertions, the saline was replaced at least once, but this was repeated more often if necessary. Then, the insertion device was pulled out (Fig. 2b, panel 4). In N=2 insertions (insertions 11 and 14), devices that had mostly been inserted were pushed in further using surgical forceps.

In N=1 rat, a second grid insertion was performed in the same craniectomy (insertions 4 & 5) to replace the previous grid, which had been mistakenly ruined by excess cyanoacrylate glue (Vetbond, cat. # B00016067, 3M, Maplewood, MN) as part of a chronic experiment. For N=2 rats, 2 insertions were performed in separate craniotomies, accounting for N=4 insertions. For N=1 experiment, the insertion was performed after a spinal cord implantation performed for another study. This rat had initially been anesthetized with ketamine. Once the spinal cord component was completed, the mote insertion continued as described above.

### Analysis of images and of videos

#### Evaluation of assembled mote grids

Once a grid was assembled, it was imaged under a benchtop microscope (SMZ645 or SMZ745, Nikon, Tokyo, Japan) using an eyepiece mounted camera (MU1803-HS, AmScope) at a variety of angles to capture multiple viewpoints. For illustrative figure making only (Fig. 3f), some images were also captured by a mobile phone camera (Pixel 6, Google, Mountain View, CA) through the eyepiece. In order to capture the lengths and angles of the carbon fibers (N=3 4x4 grids, N=6 5x5 grids, N=193 fibers), the grid was oriented such that the fibers were approximately parallel to the imaging plane and a video was captured using the AmScope camera while manually adjusting the plane of focus throughout the device. This simulated volumetric imaging commonly used with an automated stage (e.g., [72]). This process was repeated after rotating the grid approximately 90°. An image captured of a calibration slide (0.01 mm/division) (MR095, AmScope) was used to scale the videos. The two videos were loaded into ImageJ (Fiji distribution) [73] as z-stacks and a line was fit from the tip of each fiber to its base for both images. To account for the rotation of the grid in the XY plane, lines were fit to the top edges of mote bases in the closest row and these angles averaged, producing a global mote base angle against which fiber angles were compared. Fibers that were too difficult to distinguish in the videos were excluded. Once the projected length and angle of each fiber were measured in both images, basic trigonometry was performed using custom MATLAB (MathWorks, Natick, MA) scripts to determine the fiber’s length and angle in three dimensions, where the angle was defined as the deviation from the vector orthogonal to the mote base’s fiber-side.

**Figure 3.**
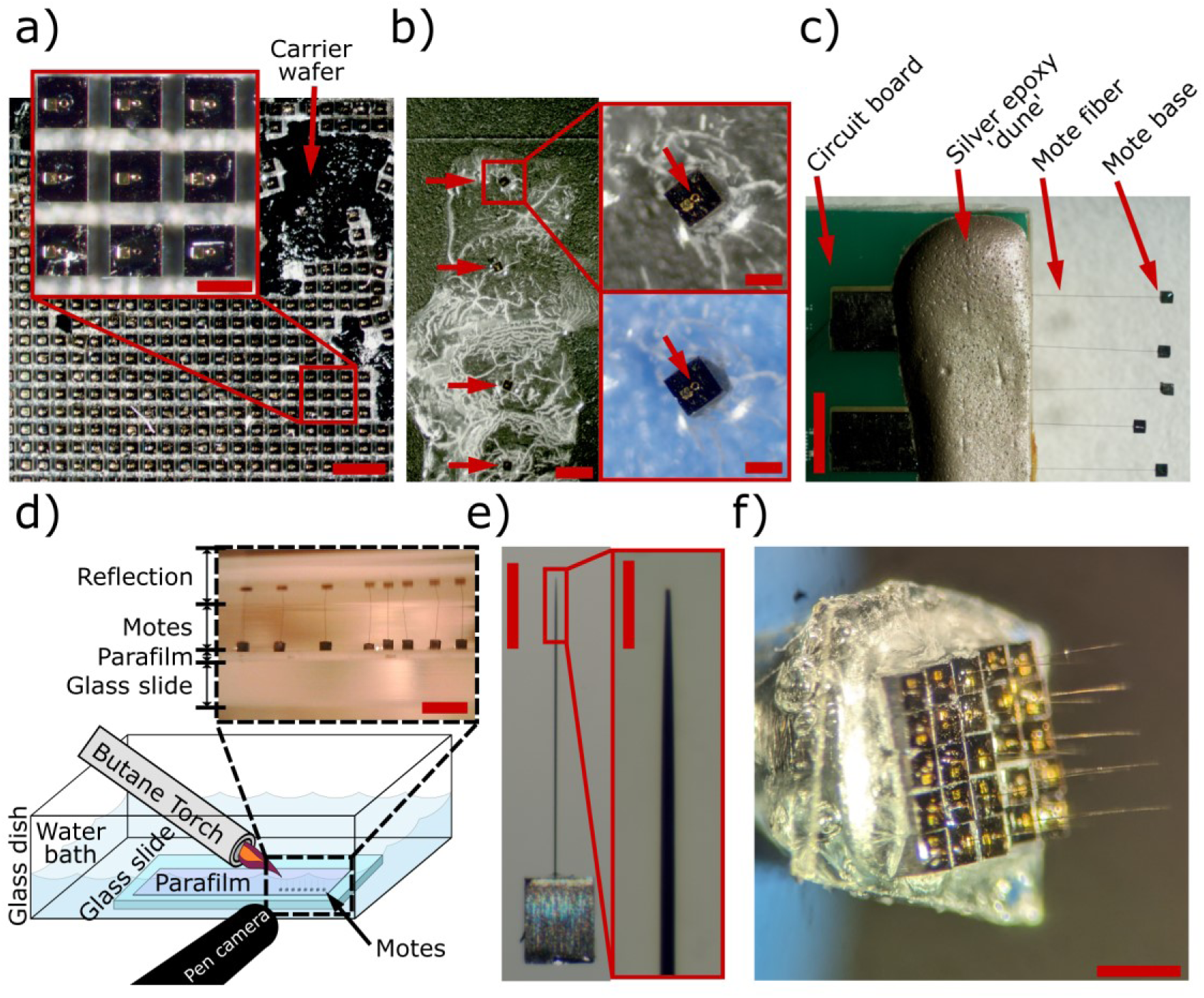
Snapshots from carbon fiber mote fabrication. a) Source non-functional mote analogs produced from cleanroom fabrication. Mote bases were attached to a thick layer of Parylene C (white) on their bottom side deposited prior to singulation. This Parylene C layer in turn was held by a carrier wafer, which was made visible after removing a subset of the mote bases (red arrow). Inset is for closer inspection of nine mote bases. Scale bar: 1 mm; scale bar (inset): 250 µm. b) During fiber placement, mote bases were held in place on a glass coverslip via PEG. Left shows a top-down view of four motes held in PEG, indicated by red arrows. Insets (right) show a closer view of a mote. To more easily illustrate the PEG (clear and white colors) for this manuscript, black electrical tape is used in the background (left, top right). During assembly, blue tape was used instead (bottom right). Arrows in the insets point to the carbon fiber holes. Scale bar (left): 1 mm; scale bars (right): 200 µm. c) Motes were suspended from a circuit board by their fibers via silver epoxy to ensure that Parylene C could be deposited on all surfaces. Image captured after Parylene C deposition. Scale bar: 2 mm. d) Fire-sharpening non-functional motes diagram (bottom) and view from pen camera (top). Motes are submerged in water to protect the fibers except at the tip when exposed to flame from a butane torch. Parafilm melted onto a glass slide is used to hold the motes while submerged since it is not water soluble, unlike PEG. The pen camera is used to monitor the water level via mote reflection so that the meniscus is touching the fiber tips during sharpening. Scale bar: 1 mm. Diagram and procedure adapted from [70]. e) Close up view of a completed non-functional mote without Parylene C encapsulation. The inset shows a close-up view of the sharpened tip. Scale bar: 300 µm. Inset scale bar: 50 µm. f) Image of a completely assembled 5x5 non-functional mote grid. The device is rotated slightly to show some bases (left) and fibers (right) in focus. Scale bar: 500 µm, estimated from mote dimensions.

Using the same microscope, AmScope camera setup, and scaling method, top-down images of completed mote grids were captured to measure the spacing between motes. One image was selected per grid and loaded into ImageJ, and quadrilaterals were fit to the fiber-side faces of the bases using the polygon selection tool. The position of a mote was defined as the centroid of the quadrilateral. The centroids were exported from ImageJ to MATLAB, and the distances between a given centroid and the centroids immediately adjacent but not diagonal to it were determined as the pitches. For example, if a mote was centrally located in the grid, four pitches were measured for that mote. Across N=6 4x4 grids and N=6 5x5 grids, 384 pitches were measured. 3.6% of these pitches were shorter than the mote horizontal side length (240 µm), which we attributed to the combination of inconsistencies in mote dimension and in the measurements themselves.

#### Evaluation of mote insertions

Images and videos collected during insertion experiments were carefully viewed to classify the insertional outcome of each mote that we attempted to insert (proof-of-concept insertions: N=45 motes, larger grid insertions: N=189 motes). These were viewed using a combination of ImageJ, VLC media player (3.0.8) (VideoLAN Org., Paris, France), and Windows (Microsoft, Redmond, WA) media viewing applications. For videos, the playback speed was adjusted and segments were viewed repeatedly. To make some video segments clearer, basic image processing techniques were sometimes employed, such as gamma correction, zoom and contrast adjustment.

Generally, a small number of motes that failed to insert were easily identified during the procedure as they either sprung away from the insertion site or fell onto their side, which is why one was immediately removed in insertion 6. For insertions with 4x4 and 5x5 grids, some motes were mostly inserted but some portion of the fiber was visibly outside of the brain. Because some of these motes were nearly entirely in the brain (e.g., motes in Fig. 7) and some were pushed in afterward with forceps (Fig. S7, Vid. 4), we made the *post hoc* decision to classify these devices as successes or failures using a threshold on the remaining fiber length. For motes where some portion of the fiber was visible (N=17), we estimated the length of visible fiber in ImageJ, where the image was scaled using nearby motes and the horizontal and vertical scales sometimes differed. If the length was ≤200 µm, the mote was classified as having been successfully inserted. Otherwise, the mote was classified as having failed to insert. We chose this threshold because the fibers had a target length of 1000 µm, and the depth where Layer V begins in rat motor cortex is 810 µm [74].

There were some cases where a small segment of the fiber was visible in the video but the mote base appeared to otherwise sit on the brain surface. These were classified as successes as this evidence pointed to those fiber segments being visible due to the transparency of the pia. Additionally, motes that were situated in inner rows and columns of the grid may have had some portion of fiber outside the brain, but side views of these devices were blocked by motes in nearby rows or columns. We classified these motes as having successful insertions as the height of their bases was equal to or less than those of motes in outer rows that had been successfully inserted. Overall, N=189 1 mm motes and N=45 preliminary motes were attempted and classified. In one insertion, N=3 of these motes were excluded because of surgeon error, where an incomplete durectomy resulted in dura blocking these devices from attempting to insert into the brain. An additional 5x5 mote grid insertion was excluded entirely because the insertion device bumped into the edge of the craniectomy when being driven toward the brain. The force of the collision was sufficiently high to shear several of the fibers such that they broke off the mote base, which had not been seen before, and was indicative that the insertion was abnormal. For proof-of-concept insertions (N=5 3x3 grids), insertion success was evaluated more stringently. Some devices that tilted with qualitatively large angles were classified as failures due to their shorter fiber length.

Videos were also carefully inspected to attribute probable failure modes that prevented motes from fully inserting via the insertion device. For this qualitative analysis, both proof-of-concept insertions and those with larger grids were considered (N=14 insertions total). First, motes were categorized into whether they had mostly inserted (N=17) with some portion of the fiber visible or had entirely failed to insert (N=15). It is important to note that motes that had mostly inserted with spans of fiber that were ≤200 µm in length and were classified as successful above were included in this analysis as motes that had mostly inserted. Devices were carefully tracked in insertion videos to determine failure modes that factored into their insertion outcome. For the majority of these motes (19/32 motes), multiple failure modes could be attributed to a given mote’s outcome, e.g., a device that started to release early but also was pushed by another mote, so all possible failure modes were attributed to each device. These failure modes were then ranked in order of estimated effect on the insertion outcome for that mote. This ranking varied in subjectivity, as some rankings could be clearly deduced from the videos while for others what could be seen in videos was more ambiguous. This ranking is why the frequencies of all identified failure modes, shown in Table S2, add up to a number greater than the total number of motes assessed (N=32). Each failure mode was then assigned to one of four primary categories: methodology, brain topography, mote interactions, or surgical complication. Each mote assessed was then assigned one of those four primary categories according to the primary category of the highest ranked failure mode, which is shown in Table 2.

**Table 2.**
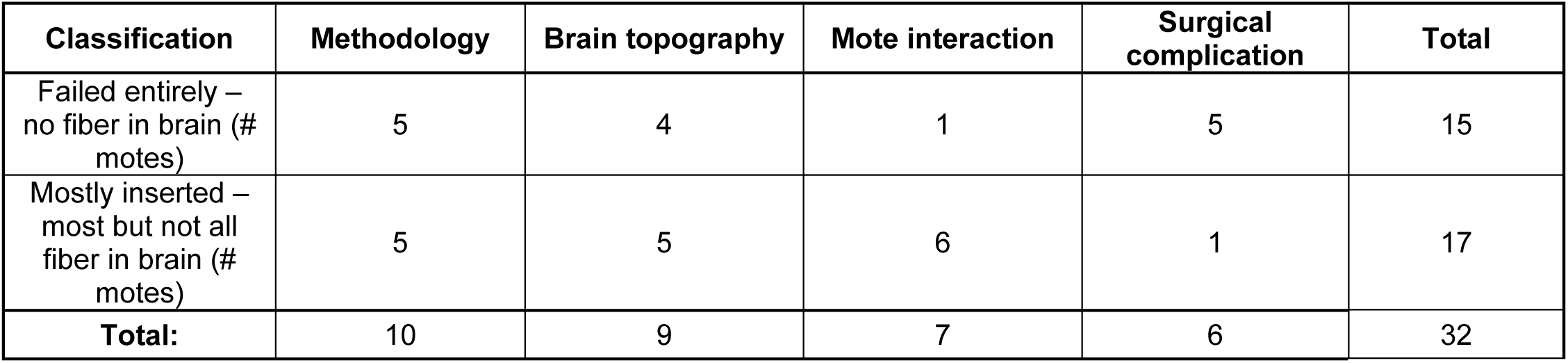
Failure modes attributed to incomplete mote insertions. Motes that failed to insert entirely (N=15 motes) and motes where the fibers mostly inserted but some portion of fiber was outside the brain (N=17) were grouped into four main categories according to the primary failure mode for incompletely inserting. Each count is the number of motes associated with that failure mode (columns) by insertional outcome (rows).

#### Mote spatial arrangement after implantation

Shortly after the removal of the insertion device, bird’s eye images were collected using the surgical scope with AmScope camera attached. Only insertion experiments where one of these photos was collected after removing the saline were used (N=5 insertions, N=104 motes). Images were loaded into ImageJ and scaled by measuring the topside edges of four motes facing the scope (the side of the mote opposite to the fiber-side) and averaging the results. Quadrilaterals were then fit to the sides visible to the scope. Motes were excluded if they had failed entirely in inserting or had topsides that were partially obstructed by other motes such that fitting a quadrilateral onto them would have been inaccurate. To measure mote pitch after insertion (Fig. 9c), the Euclidean distance between the centroid of each quadrilateral and neighboring quadrilaterals was determined using the same process as that used for pre-insertion pitch measurement.

To estimate mote displacement after insertion, pictures and insertion videos were used to determine the change in grid orientation between top-down photos captured after grid completion and the photo captured after insertion. Centroids of quadrilaterals fit for pre-insertion pitch measurements were rotated and flipped vertically to match their post-insertion arrangement. Both pre-insertion and post-insertion centroids were then centered by either the centroid of the central mote (5x5 grids) or a corner point of a centrally located device (4x4 grids). Both sets of coordinates were rotated, where the angle was defined by the slope between two mote centroids that were representative of the overall rotation of the grid post insertion. With the grids aligned, the displacement was measured as the Euclidean distance between each mote’s pre-insertion centroid position and its post insertion centroid position. Since the displacement for the center mote in 5x5 grids was 0 µm, these center motes were excluded from the final distribution (N=3 motes).

To determine mote tilt, the vertices of each quadrilateral fit to the top face were imported into MATLAB using a custom script based upon ImageJ’s region of interest code (accessible on GitHub at https://github.com/imagej/ImageJ/blob/master/ij/io/RoiDecoder.java). They were rotated according to the rotation measured during displacement estimation and translated such that the bottom left corner of the grid was at the origin. Perspective-n-point estimation [75] was performed via the *solvePnP* function of the OpenCV library [76] for each mote to determine its pose relative to the camera, where perfect top-down coordinates were used as the world coordinates. The rotation components of the poses of all motes in the insertion were averaged (Bruno Luong. RotationMean [https://www.mathworks.com/matlabcentral/fileexchange/134037-rotationmean], MATLAB Central File Exchange) for use as a reference frame, as the pose of each mote was offset since the surgical scope was not directly parallel to the surface of the brain. The tilt for each mote was calculated by determining the rotation required to rotate a mote with the reference orientation to the pose estimated by *solvePnP* and then extracting the rotation relative to the positive z-axis in the reference frame using the MATLAB *tilt* function. One insertion was excluded because the surgical camera was positioned at too large of an angle for the rotation averaging to produce a viable reference given the assumptions made about the world coordinates. Another insertion was excluded because the resultant angles were clearly incorrect, which we attribute to the poorer quality of the image. In total, N=3 insertions were used for mote tilt determination (N=73 motes).

### Statistical analyses

All statistical analyses were performed using MATLAB functions. Most statistics performed were descriptive statistics, specifically reporting the mean, standard deviation, and showing the distribution of measurements with violin plots. These were used to report and visualize the following: fiber deviation angles, fiber lengths, fiber effective lengths, mote pitch before and after insertion, deviation angle by insertion outcome, mote tilt after insertion, and mote displacement after insertion. To determine the potential relationship between fiber deviation angle and insertion outcome, multinomial and binary logistic regressions were performed using the *fitmnr* function, which also provided the χ^2^ goodness of fit. Comparing the distributions of tilt and displacement measured for motes in this study and Neurograins in Sigurdsson *et al.* [37] after implantation was performed using the following steps. Counts per bin for Neurograins were read from the histograms reported for them [37], and counts for motes were computed from our measurements but using the same binning. Then, both sets of counts were converted to proportions of the total measurements for each design and plotted together in the same histogram (Fig. 9e & Fig. 9f).

### Figures and graphics

Figures were generated using a combination of software programs. VLC media player (3.0.8) and ImageJ were used for extracting frames from captured videos. MATLAB (versions R2020b and R2024a) produced numerical plots. Violin plots were generated using a Violinplot add-on [77]. Inkscape (versions 1.1.2 and 1.3) was used to produce the figures. Videos were edited using a combination of ImageJ, Adobe Media Encoder (version 23.6.6, build 2) (Adobe Inc., San Jose, CA), and Adobe Premiere Pro (version 23.6.7, build 1) (Adobe Inc.).

### Code and data availability

Upon publication, raw data and code will be available at our lab website: https://chestekresearch.engin.umich.edu/carbon-fibers/.

## Results

### Fabrication and assembly of non-functional carbon fiber mote analogs

The primary goal of this study was to determine whether batches of motes with subcellular-scale penetrating electrodes could be implanted unassisted to cortical depths. To test our proposed implantation method and develop assembly methods for carbon fiber motes, we fabricated non-functional analogs that simulated functional devices with similar dimensions (Fig. 1), an approach used in other intracortical insertion studies [37]. Once the non-functional silicon bases were fabricated, which simulated the motes’ chips, we found that nearly all remaining assembly steps could be performed using benchtop processes (Fig. S2), as we have demonstrated previously [68,78]. A key consideration was securing the bases during assembly due to their miniscule dimensions (240-300 µm) [79]. Polyethylene glycol (PEG) proved to be a suitable choice for many of the assembly steps (Fig. 3b) except during Parylene C coating and when fire-sharpening the fiber tips. We found that hanging devices by their fibers allowed for complete coating of the devices (Fig. 3c). Heated parafilm was suitable for holding mote analogs when submerged in water for fire-sharpening (Fig. 3d). For this work, we produced at least 230 mote analogs with 1 mm target fiber length and sharpened tips (example shown in Fig. 3e) that comprised N=11 square mote grids (N=5 4x4 grids, N=6 5x5 grids) (example shown in Fig. 3f). A subset of this total (N=105 motes) were encapsulated in Parylene C. Additionally, at least 45 preliminary non-functional motes with target lengths of 0.5 mm were assembled into N=5 3x3 grids to develop the assembly and insertion protocols. From these numbers, we could reasonably conclude that our methods could produce non-functional carbon fiber motes in a sufficiently high quantity for insertion testing.

### Measuring non-functional mote assembly quality and layout

Next, we sought to measure the quality and consistency of the manual assembly process. Since carbon fiber mote chips will sit on the surface of the brain and the active site of each fiber will be at its tip [57,80], the fiber’s length and angle [60] will determine the maximum depth that the active site can reach. Also, since fibers were manually placed into the bases (Fig. S3), we expected some deviation from perpendicular. Therefore, we measured the deviation angles and lengths of fibers in completed mote grids (N=3 4x4 grids, N=6 5x5 grids) using videos taken of them through benchtop microscopes. Figure 4a shows a mote grid that had a representative distribution of fiber deviation angles along with line annotation of the closest row of motes. We note that even with the highest angular deviation of 10.1°, all 25 motes in this grid were successfully inserted into cortex (see Insertion 9 in Table 1). Fibers deviated on average only 5.7°±3.1° (X̄±S) from perpendicular (Fig. 4b) and were 1048±51 µm (X̄±S) in length (Fig. 4c). The tip’s implantation depth is expected to be 7 µm less than this total length on average due to the minimal deviation angles. As layer V in rat motor cortex spans a depth of 810-1330 µm [74], these fibers were capable of comfortably reaching the target depth intended for sampling layer V pyramidal neurons once implanted.

**Figure 4.**
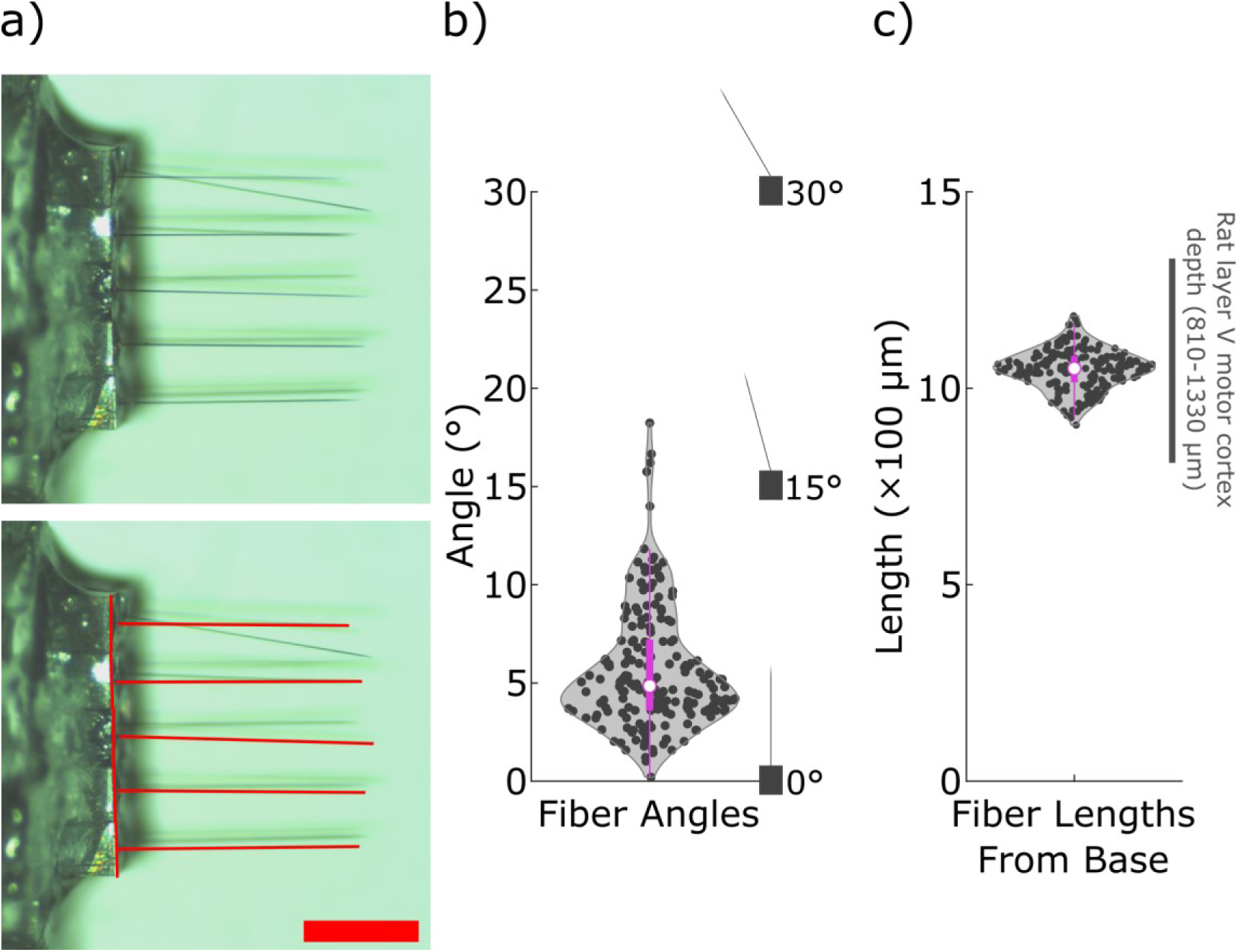
Measuring successful carbon fiber mote fabrication. a) Mote carbon fiber angles, defined as the fiber’s deviation from the vector normal to the face of the mote base from which the fiber extends, and lengths were measured from videos filmed while adjusting the focus plane throughout the grid. This process was performed twice from perpendicular viewpoints to capture perpendicular profiles of each fiber. Fitting a line to the image captured of the fiber in each viewpoint enabled calculating the fiber length and angle using trigonometry. Example frame used for measurement (top), with annotated image (bottom) that had lines fit to fibers (horizontal) and mote top faces (vertical). The latter was used to calibrate the overall angle of a given image. Scale bar: 500 µm. b) Violin plot of N=193 measured fiber angles, 5.7±3.1° (X̄±S). Cartoon profiles of motes with angles of 0°, 15° and 30° are shown for reference. c) Violin plot of measured N=193 fiber lengths, 1048±51 µm (X̄±S). The range for rat layer V motor cortex [74], the target region and depth, is shown for comparison.

We observed early on that the devices readily aggregated together when partially submerged in the PEG used to affix them to the insertion device. This is shown in Video 1 in two ways: first, when a mote base placed onto an insertion device is pushed into heated PEG, it is drawn toward the other bases; and second, the grouped mote bases remain aggregated when pushed by forceps in heated PEG. This phenomenon minimizes extra space between motes and helps us realize an electrode density close to the theoretical maximum based on mote dimensions. We therefore prepared all grids in this study with motes aggregated as closely as possible (Fig. 3f, Fig. 5a). To determine the resulting pitch, we captured top-down images of mote grids (N=12 grids, N=6 4x4 grids and N=6 5x5 grids) and measured the center-center pitch from the fiber-sides that were visible (Fig. 5a). The resulting pitch (251±8 µm, N=384 distances) was only 11 µm larger on average than the motes’ nominal side lengths (240 µm) (Fig. 5b).

**Figure 5.**
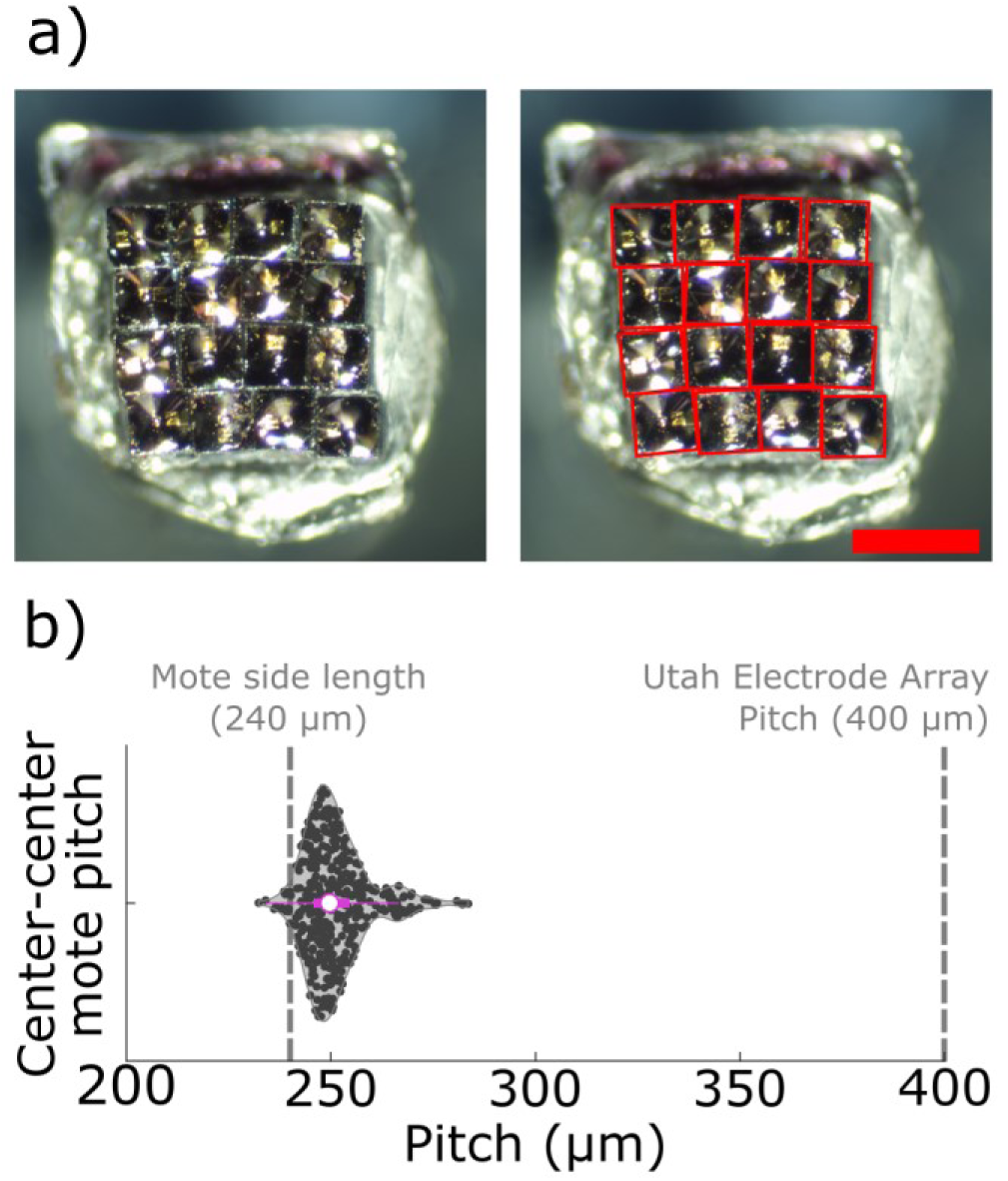
Measuring mote grid spacing after assembly. a) The pitch of a completed grid was measured by capturing a top-down image of the grid, fitting quadrilateral ROIs to the top faces, and calculating the distances between quadrilateral centroids. A representative image (top) with fit quadrilaterals shown in red (bottom). Scale bar: 500 µm. b) Violin plot of N=384 pitches measured from N=12 grids (N=6 4x4s, N=6 5x5s), 251±8 µm (X̄±S). The mote side length (240 µm) and UEA electrode pitch [81] are shown for comparison, where the former is the possible pitch minimum and the latter is for comparison against the gold standard.

**Figure 6.**
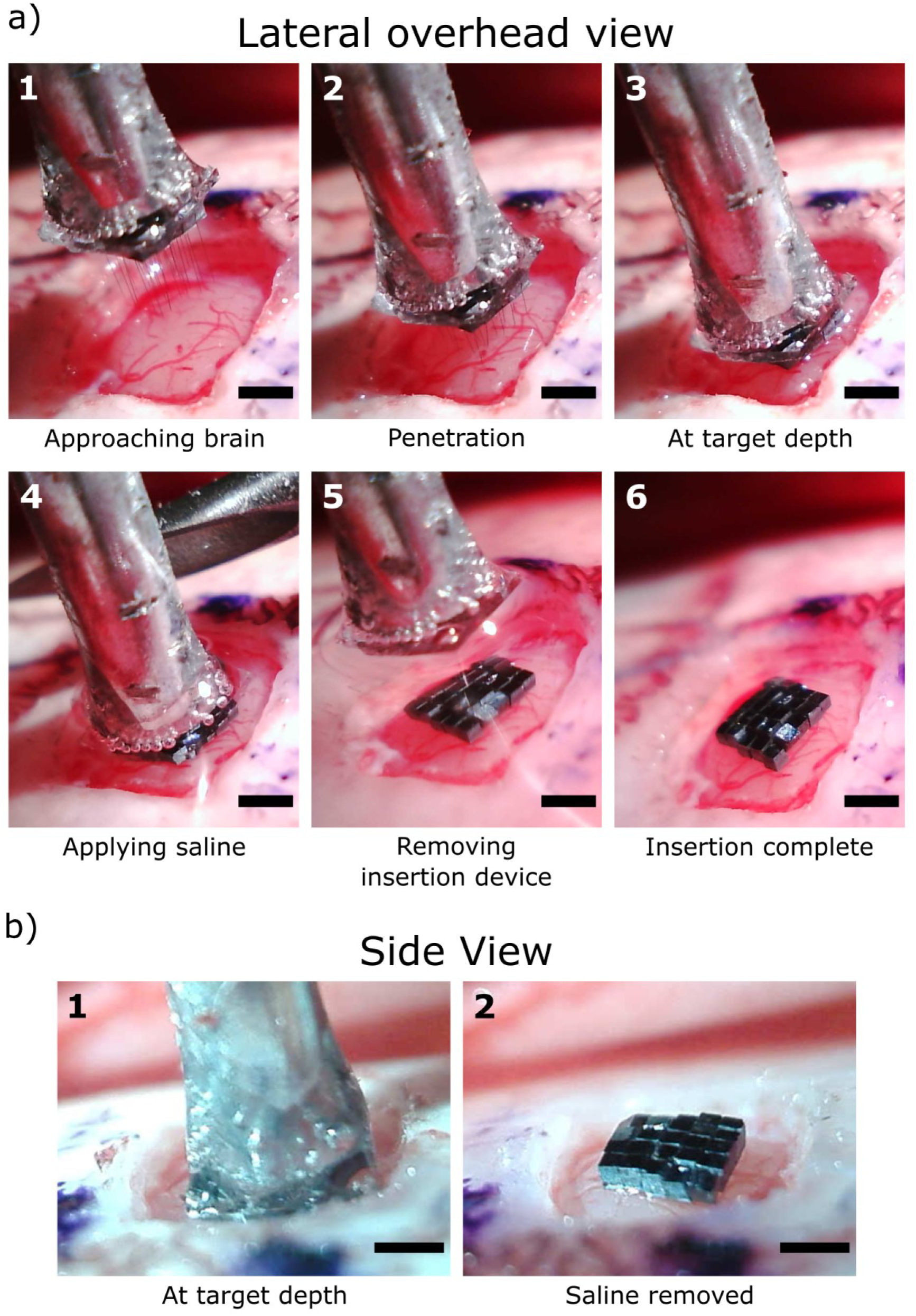
Insertion of non-functional carbon fiber motes into rat cortex. Frames extracted from videos captured during an exemplary implantation of a 5x5 mote grid where all 25 motes successfully implanted. a) Frames captured from a lateral overhead view. The order of the frames is shown numerically in the top left corner of each frame. See Vid. 2 for source footage. Scale bar: 1 mm. b) Frames extracted from a side view approximately level with the craniectomy. See Vid. 3 for source footage. Scale bar: 600 µm.

**Figure 7.**
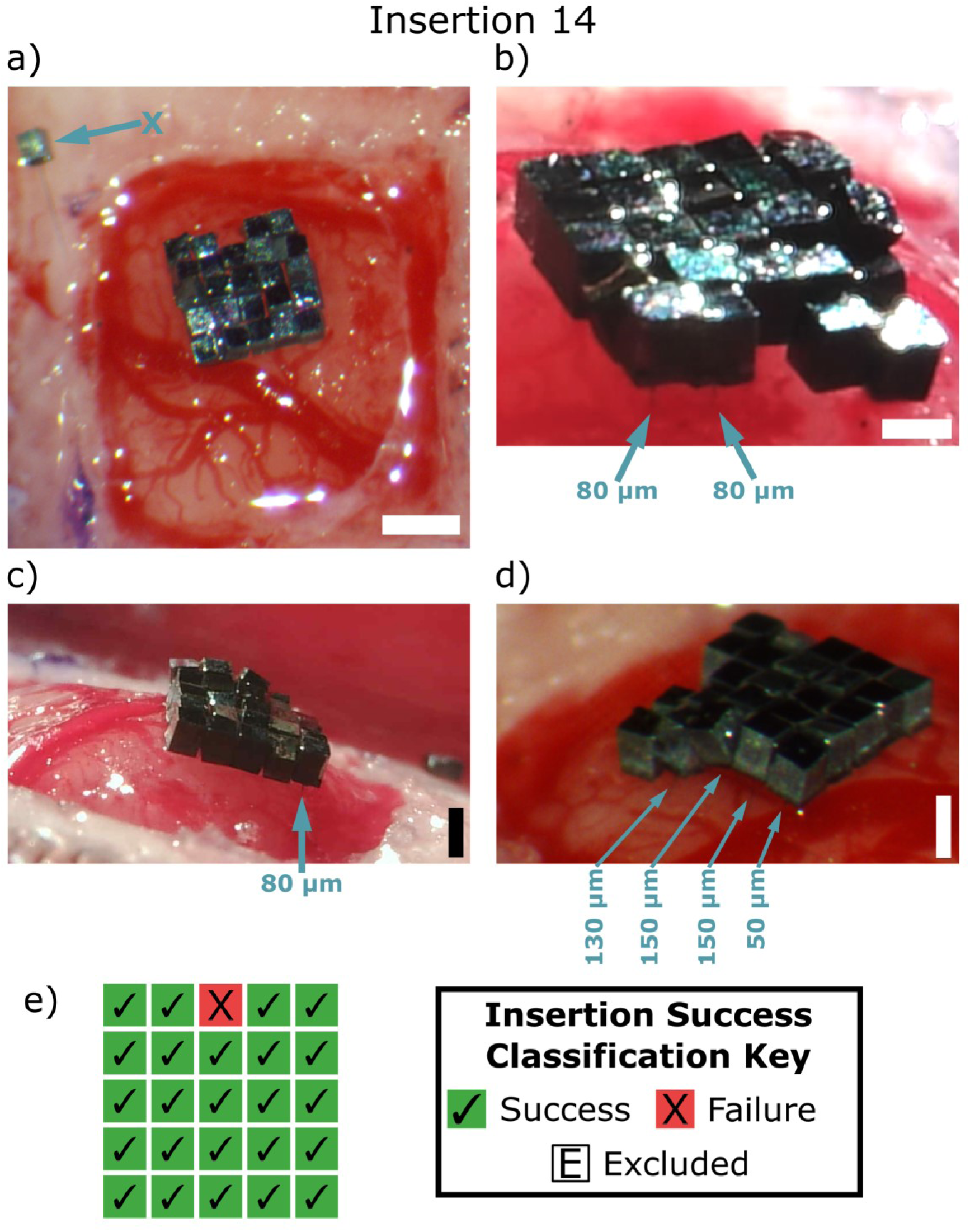
Illustrative example of classifying an insertion with representative outcomes. Multiple viewpoints of an insertion shortly after removal of the insertion device: a) bird’s eye view, scale bar: 750 µm; b) side view from the anterior edge of the craniectomy, scale bar: 300 µm; c) side view from the medial (right) side of the craniectomy, scale bar: 400 µm; d) lateral & posterior overhead (vertically from the bottom left corner of the craniectomy) view, scale bar: 400 µm. Motes that successfully inserted with 200 µm or less of fiber remaining (N=6) are labelled with arrows and the estimated length of fiber that is still outside of the brain, rounded to the nearest 10 µm. The mote that entirely failed to insert (N=1) is labelled with an arrow and the letter “X”. e) Insertion outcome classification diagram for this insertion. The layout matches that shown in the bird’s eye view. Image processing: a) none; b) 10 frames of video captured were averaged, then gamma corrected and contrast adjusted in ImageJ [73]; c) none; d) gamma corrected in ImageJ.

### Successful insertion of non-functional carbon fiber mote analogs

With mote analogs successfully assembled, we sought to evaluate how they, and by extension our proposed functional designs, could be implanted into the brain. Using priorities defined by the literature, we aimed to design an implantation procedure with the following attributes: 1) efficient, i.e. inserting tens to hundreds of electrodes simultaneously [37]; 2) rapid, in order to avoid adding an unnecessary time burden on the patient and surgeons [27,82]; 3) facile, in order to avoid further complicating neurosurgical implantation [27,82]; and 4) minimally invasive to reduce insertional trauma [83]. The resulting implantation procedure used PEG to hold devices onto an insertion device [37,38,84] while the fibers penetrated the brain and mote bases were lowered to the surface. Once in place, the devices were released via dissolution of the PEG after exposure to saline and body heat over the course of seconds to minutes (Fig. 2). We validated this method *in vivo* by implanting square mote grids (N=14) of multiple configurations into rat cortex (Table 1 and Table S1). To illustrate the insertion procedure, we show video frames captured during the insertion of a 5x5 non-functional mote grid when all 25 devices were successfully implanted. Figure 6a and Video 2 show the procedure from an overhead point-of-view, whereas Figure 6b and Video 3 show the procedure from a viewpoint level with the craniectomy. As these image sequences demonstrate, the carbon fibers could penetrate the brain during insertion and would retain a nearly square configuration once the insertion device and saline were removed.

As each procedure entailed the implantation of multiple motes simultaneously, we defined the success rate for an insertion test as the sum of outcomes observed for each mote attempted. For proof-of-concept insertions using preliminary devices (N=5), we implanted grids with low numbers of motes (3x3 grids) that had short fibers (∼500 µm) and blunt tips. With 39 of 45 (87%) attempted preliminary motes implanting successfully (Fig. S4 & Table S1), these initial successes motivated increasing the grid size and fiber length in subsequent insertions (N=9) to more closely simulate functional device configurations. Overall, larger grids consisting of 16 and 25 motes and 1 mm long fibers with sharpened tips also successfully inserted into the brain using this implantation method. Across N=9 insertions (N=4 4x4 grids, N=5 5x5 grids), 171 out of 186 motes were inserted successfully for an overall success rate of 92% (Table 1). Bird’s eye views of these insertions along with diagrams of their insertion classifications are shown in Figures 7, S5 and S6. In particular, Figure 7 illustrates the classification of a successful grid insertion that was representative of most insertions, and Figure S5 shows classification of an insertion with poor outcomes.

### Characterizing failure modes observed during carbon fiber mote insertion

Although the success rate for mote implantation was high, we identified several failure modes during these insertions to determine mitigation measures for future implants. Using the videos and images collected during implantation surgeries (N= 14 insertions, 231 motes), we first identified motes that failed entirely to insert (N=15) or mostly inserted but still had a portion of the fiber visible outside of the brain (N=17). We make this distinction because in two experiments, we pushed in motes that had mostly inserted by simply nudging them with forceps, as shown in Vid. 4 and Fig. S7. Next, we examined these attempts more closely using surgical videos and notes. The full list of putative failure modes and their frequencies is presented in Table S2, while counts according to overarching categories for these failure modes are shown in Table 2. Attributes of our assembly and insertion methodology accounted for the highest proportion (31%) of incomplete mote insertions, with buckling and early PEG release as common failure modes. Brain topography was the second highest in proportion (28%), where dimpling from large blood vessels and swelling were the most common. These were followed by mote interactions (22%), and surgical complications (19%). With the exception of mote interactions, which were difficult to predict, acting on failure modes associated with the other three categories improved outcomes as insertion experiments were performed, such as removing the insertion device with a vertical motion or more vigilantly removing blood.

We next investigated whether variation in carbon fiber angle from perpendicular, which approximated the fiber angle relative to the cortical surface, could negatively influence insertion success [85]. Therefore, we explored how the fiber deviation angles measured after grid completion (Fig. 4a-b) might have factored into insertional outcomes. Figure 8 shows the deviation angles for motes in N=7 mote grids (N=154 total motes) categorized by insertion success. Interestingly, the angles associated with motes that failed entirely (5.9±4.2°, X̄±S N=8 motes) and those that mostly inserted (6.7±4.6°, N=16 motes) were comparable to those that fully inserted (5.3±2.5° X̄±S, N=130 motes). Moreover, when we used multinomial logistic regression to determine whether deviation angle might be a factor in insertion outcome, we found that the angle was a nonsignificant predictor (p=0.2, χ^2^ goodness of fit) at these small fiber deviations (< 16°). Fiber angle was still nonsignificant even when failed and mostly inserted motes were pooled (p=0.09, χ^2^ goodness of fit). Overall, these results suggest that fiber angle may not be the primary factor in insertion outcome given the deviation angles in this study.

**Figure 8.**
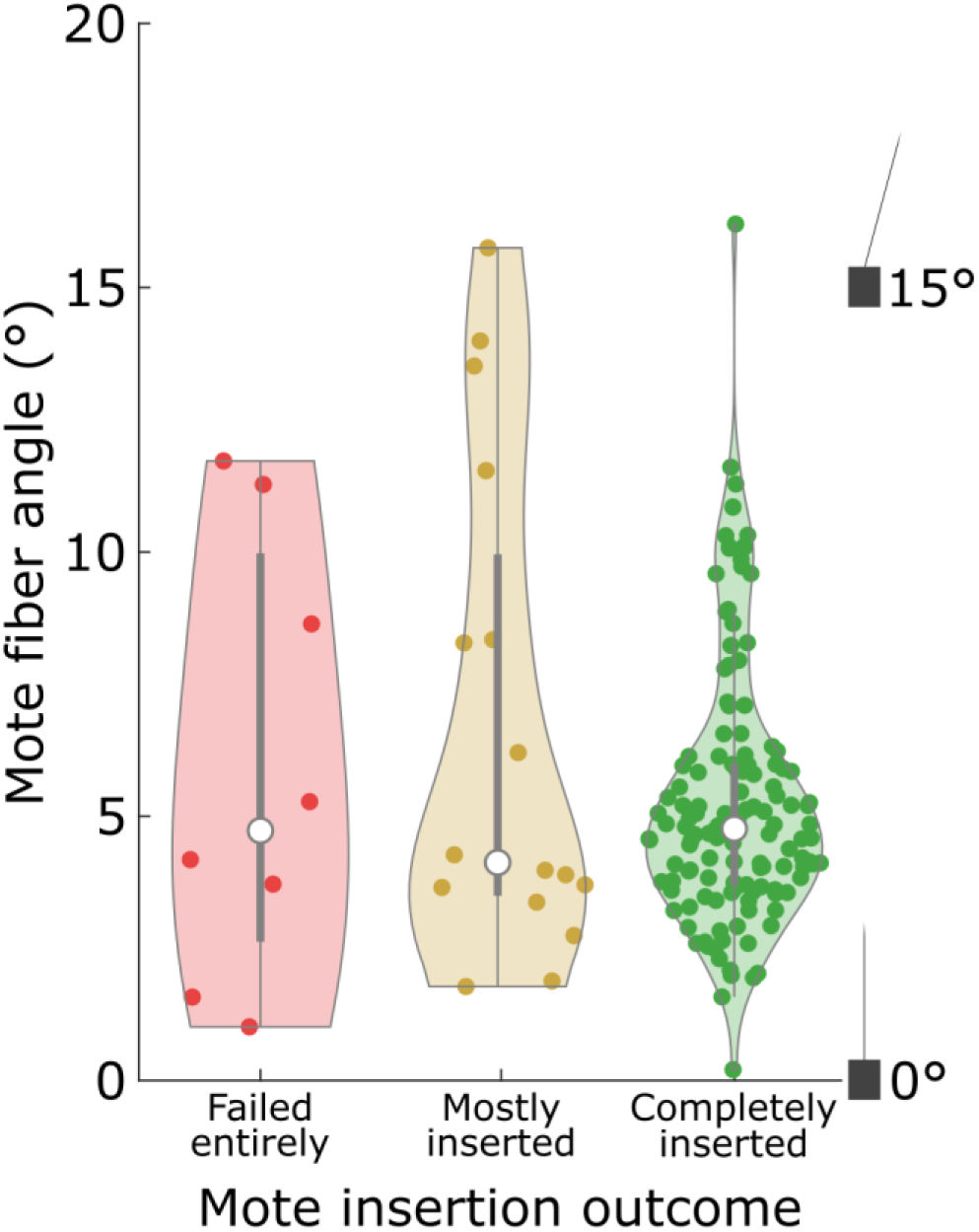
Mote fiber angle categorized by insertion outcome. The fiber angles that were measured after grid completion and prior to implantation (see Fig. 4a-b) for N=7 mote grids were categorized by the outcome of attempting to insert the mote associated with each angle measurement (N=154 angles). As in Figure 4a-b, the mote fiber angle here is defined as the deviation from the vector normal to the face of the mote base from which the fiber extends. Angles for motes that failed to insert: 5.9±4.2° (X̄±S), N=8. Angles for motes that mostly inserted: 6.7±4.6° (X̄±S), N=16. Angles for motes that successfully and completely inserted 5.3±2.5° (X̄±S), N=130. Motes that were excluded from insertion outcome classification are not shown (N=3).

### Device arrangement after implant

Since NIR communication is “line of sight” by definition, where a mote’s topside points may affect its capacity to harvest power and communicate with the repeater unit [22,63]. Therefore, we calculated motes’ arrangement immediately after implantation by measuring their changes in position (Fig. 9a) and tilt from horizontal (Fig. 9b), metrics defined previously in the literature [37,38]. To do so, we collected bird’s eye photos of motes situated on the brain’s surface and manually fit quadrilaterals to the topsides of motes that had fully or mostly inserted (N=3 5x5 grid insertions, N=2 4x4 insertions, N=104 motes). The resulting pitch was 281±43 µm (X̄±S, N=153 pitches), a 12% (31 µm) increase in average pitch from the 251±8 µm (N=168 pitches) measured after grid assembly (Fig. 9c). We used a subset (N=3 5x5 insertions, N=73 motes) of these quadrilaterals to determine the tilt angles for implanted motes, which averaged 22±9° (X̄±S) as shown in Figure 9d. Figure S8 shows the annotated source images with corresponding angles for easier visualization of the tilt measurement. To compare these results to prior work, we plot mote displacement (65±55 µm, X̄±S, N= 101 motes) and tilt against similar measurements reported for Neurograin chips acutely implanted into rat cortex [37] in Figures 9e and 9f, respectively. These histograms show that the distributions for mote displacement and tilt measurements skewed lower compared to those measured for Neurograins. Similarly, the percentage of carbon fiber motes that were displaced more than 200 µm was only 5% and the percentage that tilted more than 45° was only 3%, which were lower than the proportions observed for Microbead chips acutely implanted into rat cortex (9% and 36%, respectively) [38]. Overall, these results suggest that carbon fiber motes may retain their arrangement throughout the implantation procedure to a greater degree than other chips that are implanted inside the brain.

**Figure 9.**
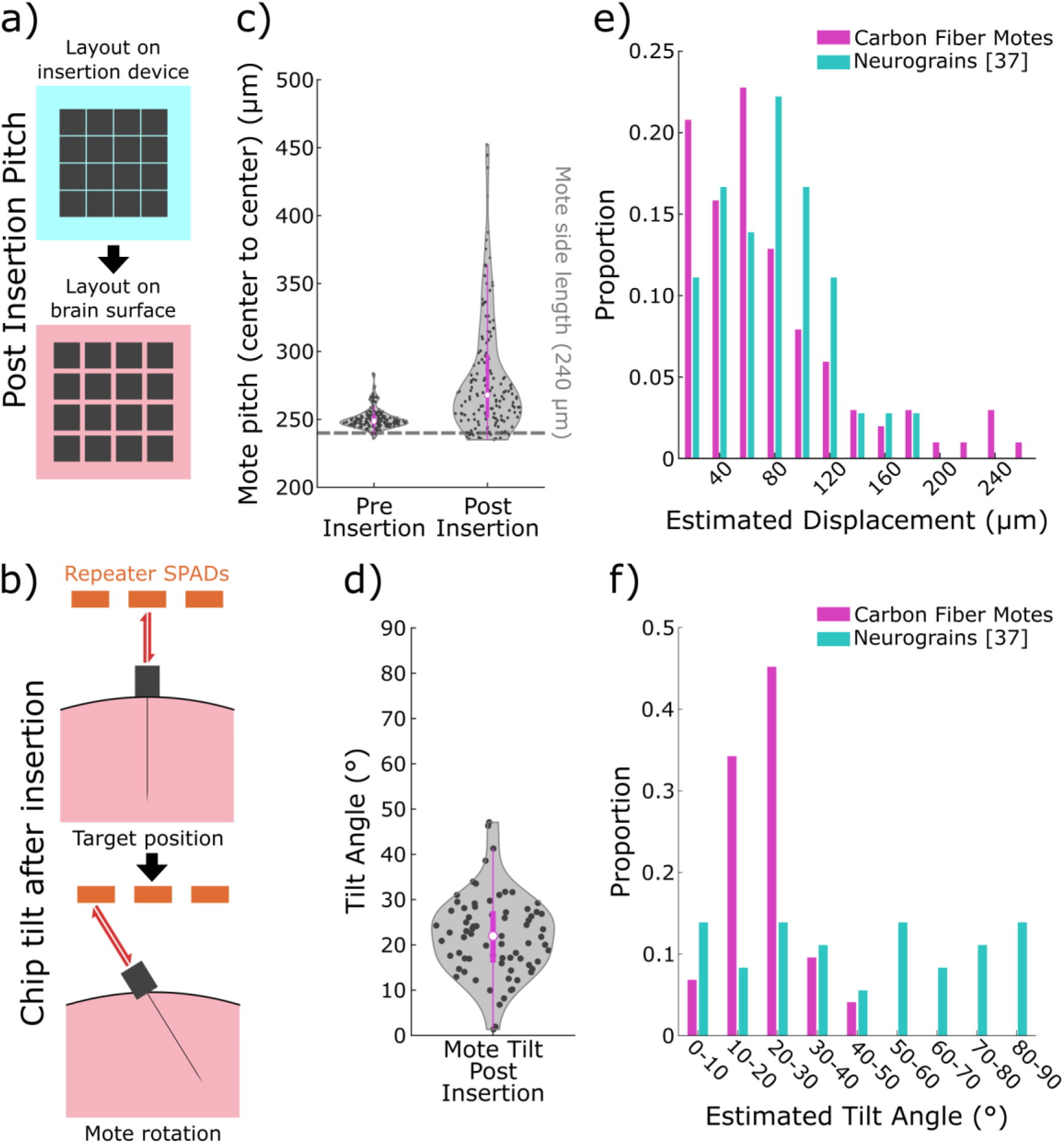
Carbon fiber mote spatial arrangement immediately after implantation. a) Cartoon illustrating an increase in the pitch between motes after implantation (bottom) relative to pitch achieved on the insertion device (top). b) Cartoon illustrating how motes tilting on the brain surface can affect power harvesting and communication with the repeater unit. Top: an ideally implanted mote with no tilt communicates with the repeater directly above it as intended. Bottom: more realistically, after implantation a mote will tilt to some degree, which could point to other single photon avalanche diode (SPAD) detectors. Orange rectangles: SPAD detectors. Red arrows: near infrared light power downlink and data uplink. Pink: brain. c) Pitch measurements for N=5 mote grids before insertion (251±8 µm, X̄±S, N=168 pitches) and after insertion (281±43 µm, N=153 pitches). The pitch increased by 12% on average. The mote side length of 240 µm is shown for comparison. d) Tilt angles measured for carbon fiber mote analogs implanted onto the surface of the brain were 22±9° (X̄±S, N=3 insertions, N=73 motes). e) Comparison to displacement measured for non-functional Neurograin analogs implanted intracortically into rat cortex in acute conditions. Values for Neurograins were read off of the histogram published by Sigurdsson *et al.* [37]. The values for carbon fiber motes were estimated by calculating the change in position (65±55 µm, X̄±S, N=101 motes) after aligning pre and post implant images. These values were separated into 20 µm bins for comparison. Both distributions of histogram counts were converted to proportions for a direct comparison. f) Comparison to tilt angles measured for non-functional Neurograin analogs implanted intracortically into rat cortex in acute conditions. Values for Neurograins were read off of the histogram published by Sigurdsson *et al.* [37]. The values for carbon fiber motes are the same as shown in d), but separated into 10° bins. Both distributions of histogram counts were converted to proportions for direct comparison.

## Discussion

Neural dust offer a solution to the hardware challenges facing the conventional wired intracortical arrays currently used in human BMIs, namely safely accommodating many recording channels [86]. However, current dust chips are still orders of magnitude larger than neurons, and therefore intracortical placement of the entire dust chips may cause an adverse foreign body response (FBR) and displace large quantities of neurons [21,39,40]. Here, we evaluated whether neural dust could be implanted in batches using an approach that is less disruptive to the brain. In particular, we assembled and implanted nonfunctional motes that sat on the brain surface and could reach target cortical depths with penetrating carbon fiber electrodes. Our implantation method could implant up to 25 motes into rat cortex quickly and simultaneously with an overall success rate of 92% (171 of 186 attempted motes) such that only the 6.8 µm carbon fiber electrode occupied brain volume.

Despite the potential advantages of this approach, we found that we needed to introduce a method for implanting batches of motes onto the cortical surface due to a lack of batch methods in the literature. Whereas previous work has demonstrated procedures for implanting chips inside the brain in batches [37] or singly in rapid succession [38], more recent neural dust work has instead opted for forceps placement of chips onto the brain one-by-one [46,47]. This shift is likely due to the FBR observed from implanting devices into the brain directly [37,44]. An important benefit of our implantation method was the ease and lack of complexity involved for the surgeon. We showed that each batch of carbon fiber motes secured by PEG could be implanted in a simple micromanipulator movement toward the brain, followed by a simple removal of the insertion device after the PEG dissolved. Combined with the ability of sharpened carbon fibers to easily penetrate the brain surface [61,70], including the leptomeninges, our insertion method is simple and does not rely on specific insertion speeds or specialized equipment, like the pneumatic inserter required for UEAs [82], nor does it require extraneous technical ability of the surgeon. Another advantage is compatibility with conventional micromanipulators, and therefore the possibility of translation to several similar surgical methods used in humans. This includes the use of a mechanical arm, frame-based stereotaxy, or a neurosurgical robot [87,88]. Furthermore, adjunctive neuronavigation is possible, utilizing preoperative magnetic resonance imaging (MRI) to plan one or multiple implant locations [89]. For example, 10-20 stereoelectroencephalography electrodes can be placed in one surgery, taking approximately 5 minutes per electrode with robotic-assisted trajectories guided by preoperative imaging [88]. Avoiding contributing complexity to neurosurgery is critical in enabling the translation and clinical adoption of neural dust [27], and the ease of insertion demonstrated here promotes the use of carbon fibers in other neural dust designs.

Restorative BMIs will require neural interfaces that not only have a high recording yield but can also sample neural activity with temporal stability [90]. Previous work has linked diminished performance with daily signal instability [91] and a gradual reduction in signal quality over time [92]. These reductions in efficacy have classically been attributed to the FBR [93], although improved signal acquisition may not necessarily accompany an improved tissue response [94]. Regardless, implanted carbon fiber arrays are associated with both a minimal reaction, including minimal glial activity and the retention of nearby neurons [69,71], and a higher recording yield than silicon shank probes [69]. Since we did not record electrophysiology in this study and there is a lack of literature that includes the implantation of networks of dust-like probes, it is difficult to predict how carbon fiber motes might fare in recording yield compared to other neural probes and neural dust. However, some predictions can be made based upon the study of UEA longevity and other studies utilizing carbon fibers. In one such effort, the majority of UEAs (33/55) had maximum recording yields that were less than 80%, which further decreased over time [95]. Conversely, virtually all carbon fiber electrodes (97%) were reported to have recorded large spikes (>100 µV_peak-peak_) intraoperatively in one study [80], which suggests that they can start with higher recording yields. Moreover, the proportion of channels yielding spikes for laboratory-assembled carbon fiber electrode arrays remained an order of magnitude higher than for Michigan style probes over approximately three months [69]. With the improvements inherent in the production of medical implants and optimizing mote insertion beyond the 92% success rate reported here, we estimate that a network of wireless carbon fiber motes may be capable of fulfilling the need for high and stable recording capacity in restorative BMIs.

Although the success rate when inserting up to 25 motes simultaneously was high, achieving the thousands of channels that may be required for high-performance BMIs [23] may require some improvements to the implant method reported here. Behind improving the consistency of assembly (“Methodology” column in Table 2), we found that brain topography was putatively the most common failure mode for mote insertion. This suggests that brain surface geometry is a principal determinant in insertion success with epicortical dust devices. Here, we implanted into rat cortex, which lacks the gyral complexity of humans and other large mammals [96,97]. When implanting into the latter, the number of motes that can be delivered simultaneously will likely be delimited by sulcal boundaries. Furthermore, the greater number of penetrating electrodes may increase the “bed of nails” effect, which can become prohibitive in perforating the pia and escalate insertional trauma from the ensuing compression of dimpling [98,99]. Several methods have been proposed to diminish this effect by facilitating pial penetration [99], including pia removal [100], thinning the pia by laser ablation [99], stiffening the pia [101], sharpening electrode tips [83,102], increasing insertion speed [82] or simply increasing electrode spacing [103]. More basic research is required to determine which techniques can reduce disruption the most, however, as even well-controlled insertion experiments have shown that tradeoffs in reducing bleeding and cell disruption can render the best course of action inconclusive [102]. Since the electrodes used here had sharpened tips and we still observed dimpling, a combination of techniques may be required. A solution made possible by the modular nature of motes could be implanting several small batches sequentially, as has been previously shown with NET probes [53], and is a viable strategy in humans.

A limitation of our approach was that motes were assembled entirely by hand after the ‘chips’ were batch fabricated in the cleanroom. Although the inherent variability introduced by manual placement did not diminish the fiber lengths such that the recording sites would be outside the target cortical layer, larger fiber deviations did appear to be detrimental to insertion success in some cases, even though these angles were sufficiently small to be a nonsignificant factor overall. A more repeatable process akin to the automated carbon fiber assembly systems reported previously [104,105] could reduce the variability in carbon fiber angles considerably. For example, fibers placed robotically into micromachined shanks had deviation angles that were an order of magnitude smaller than those presented here [105]. Manual assembly also rendered the process more tedious and time consuming, where transferring motes between the substrates that secured them expended the most time. While holding devices in PEG during assembly offered a high degree of freedom, its use precluded gentler handling methods, e.g., vacuum pickup [46], and its irregularity would hinder implementing an automated or semi-automated process. Using a regularly-spaced mold or substrate formed around the mote chips could improve the regularity of the process and its efficiency, as has been shown when processing Neurograins in a cleanroom [79]. In any case, we built upon our existing benchtop assembly methods that have produced successful carbon fiber probes in prior work [78] as a baseline. That assembling wireless mote analogs with benchtop-compatible methods was possible at all conjures up the possibility of similar open-source fabrication using chips ordered from a semiconductor manufacturer.

## Conclusion

In this study, we designed and tested a simple and reliable method that could simultaneously implant up to 25 motes into the brain with an overall success rate of 92% and favorably low tilt and displacement after insertion. While increasing the array size and translation into large mammals and human brain may bring additional challenges, ours is a viable method for neural dust placement that, unlike other conventional intracortical arrays, does not require specialist skill or equipment and may be inserted with existing tools used in human neurosurgery.

## Acknowledgements

We would like to thank Khalil Najafi for allowing use of his equipment for fabrication. We also thank the staff of the Lurie Nanofabrication Facility for their expertise. We also thank Jeanpaul Posso for help with Parylene C deposition, Chris Andrews for help with statistics, Eric Kennedy for surgical assistance, and Scott F. Lempka for comments on the manuscript. This work was financially supported by the National Institute of Neurological Disorders and Stroke (UF1NS107659, RF1NS128667, & R01NS118606), the National Institute of Mental Health (1RF1MH120005-01) and the National Science Foundation (1707316 & 2129817).

## CRediT Author Statement

Conceptualization: JGL, MB, JDW, HK, DB, CAC. Data Curation: JGL, JLWL, MGC, CAC. Formal Analysis: JGL, JLWL, MGC, CAC. Funding Acquisition: DC, JDW, DB, CAC. Investigation: JGL, JLWL, MGC, MB, PRP. Methodology: JGL, JLWL, MGC, PRP, JMR, JL, JP, DB, CAC. Project Administration: HK, JDW, JP, DB, CAC. Resources: DC, JDW, JP, DB, CAC. Software: JGL, MGC. Supervision: JDW, JP, DB, CAC. Validation: JGL, JLWL, MGC. Visualization: JGL, JL, JDW, CAC. Writing – Original Draft: JGL, JLWL, MGC, MB, CAC. Writing – Review & Editing: All authors.

## Ethics Statement

All animal procedures were approved by the University of Michigan Institutional Animal Care and Use Committee.

