## Supplementary material & video legends for "A method for efficient, rapid, and minimally invasive implantation of individual non-functional motes with penetrating subcellular-diameter carbon fiber electrodes into rat cortex"

Joseph G. Letner[1,2], Jordan L. W. Lam[3], Miranda G. Copenhaver[1,2], Michael Barrow[4], Paras R. Patel[1,2], Julianna M. Richie[1,2], Jungho Lee[4], Hun-Seok Kim[4], Dawen Cai[5,6,7], James D. Weiland[1,2,8], Jamie Phillips[9], David Blaauw[4], Cynthia A. Chestek[1,2,4,6,10,*]

[1]Department of Biomedical Engineering, University of Michigan, Ann Arbor, MI, 48109, USA
[2]Biointerfaces Institute, University of Michigan, Ann Arbor, MI, 48109, USA
[3]Department of Neurosurgery, University of Michigan, Ann Arbor, MI, 48109, USA
[4]Department of Electrical Engineering and Computer Science, University of Michigan, Ann Arbor, MI, 48109, USA
[5]Department of Cell and Developmental Biology, University of Michigan Medical School, Ann Arbor, MI, 48109, USA
[6]Neuroscience Graduate Program, University of Michigan, Ann Arbor, MI, 48109, USA
[7]Biophysics Program, University of Michigan, Ann Arbor, MI, 48109, USA
[8]Department of Ophthalmology and Visual Sciences, Kellogg Eye Center, University of Michigan, Ann Arbor, MI, 48105, USA
[9]Department of Electrical and Computer Engineering, University of Delaware, Newark, DE, 19716, USA
[10]Robotics Department, University of Michigan, Ann Arbor, MI, 48109, USA

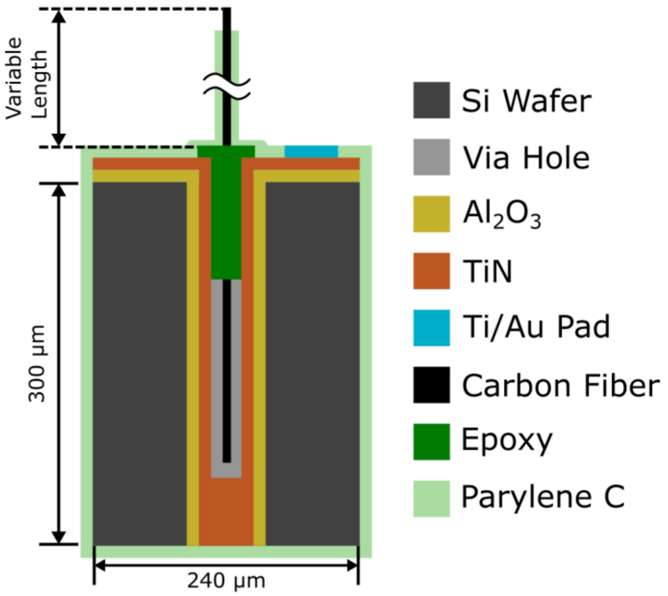


**Figure S1. Cross-sectional diagram of non-functional mote analogs.** Non-functional motes were fabricated with conductive holes. For this work, only insertion testing was performed, so the conductive capabilities were not used. Also, many devices were not encapsulated in Parylene C for insertion testing (see Table 1 and Table S1).

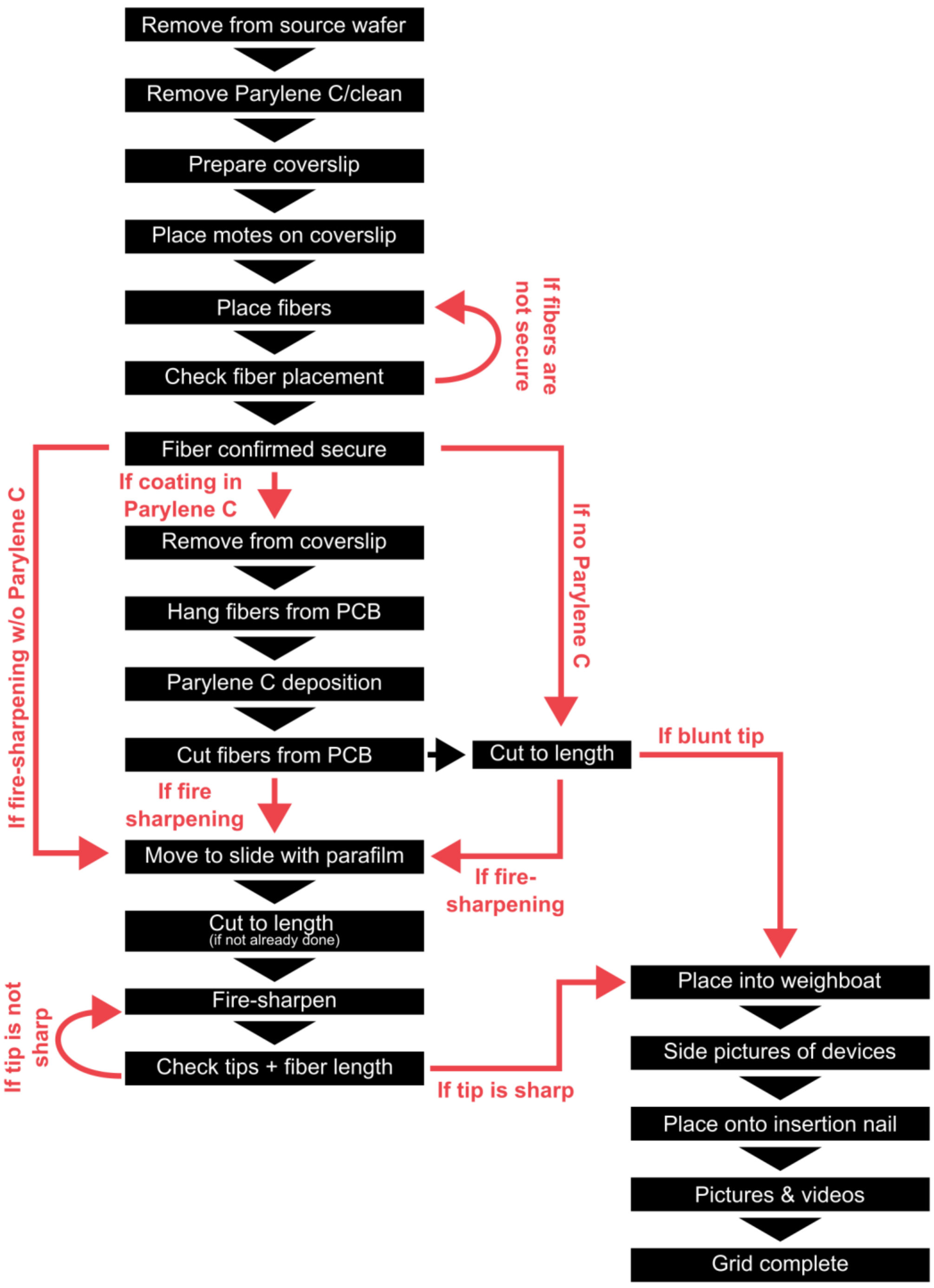


**Figure S2. Assembly steps flow for non-functional carbon fiber mote analogs.** This diagram shows the flow of steps used for assembling non-functional carbon fiber mote analogs after the silicon bases were micromachined in a clean room. All steps are benchtop processes, with the exception of Parylene deposition. See the methods text for more details. Four different types of motes were used in this work: 1) preliminary motes with target lengths of 500 μm and blunt tips, used for proof-of-concept insertions (N=5 insertions), 2) motes with target lengths of 1 mm and sharp tips (N= 5), 3) motes with target lengths of 1 mm and sharp tips, where all but the tip was coated in Parylene C (N=4), and 4) motes with target lengths of 1 mm and sharp tips reused from previous insertions (N=1). The steps above cover the steps for the first three cases (reused motes were simply cleaned), which is why there are multiple pathways to a completed grid of motes. Light red colored arrows and text denote steps that could differ depending upon the type of mote being assembled. PCB = printed circuit board.

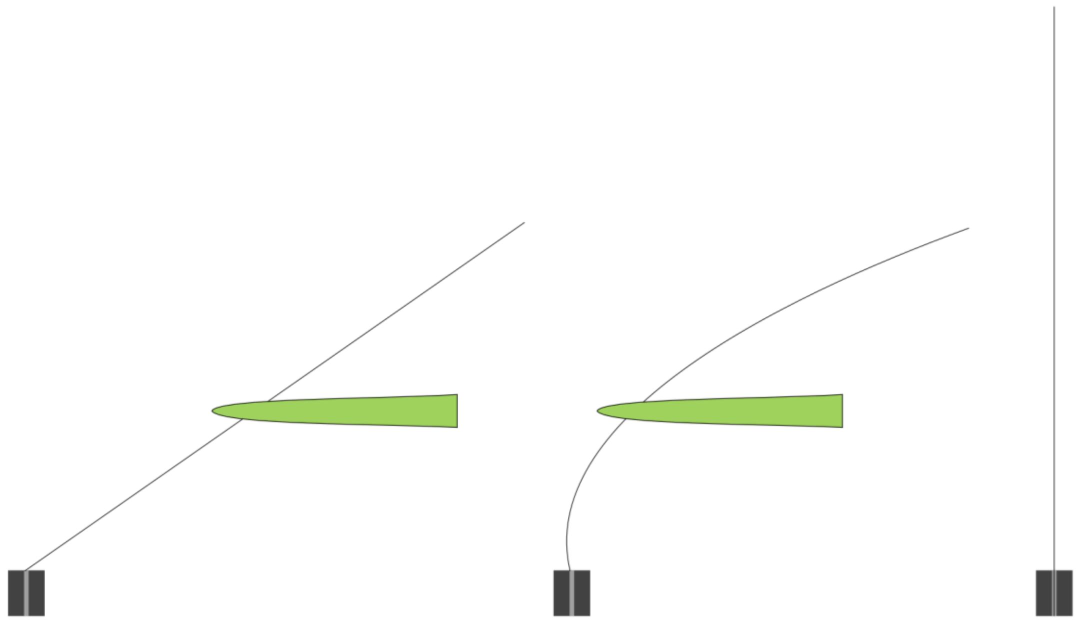

**Figure S3. Threading carbon fibers into mote base holes.** Shown here is a three-part action sequence illustrating the motions generally used to thread carbon fibers into the holes of carbon fiber mote bases. First, the fiber is brought to the hole with forceps with a lateral orientation. Once the end of the fiber is at the top of the hole, the fiber is pushed downward and laterally to force the fiber to bend. Once enough tension is applied to the fiber, it springs downward into the hole and the fiber is release from the forceps. Green curved structure: fine tipped forceps. Black line: carbon fiber. Dark grey rectangle: mote fiber base. Light rectangle: hole in mote base.

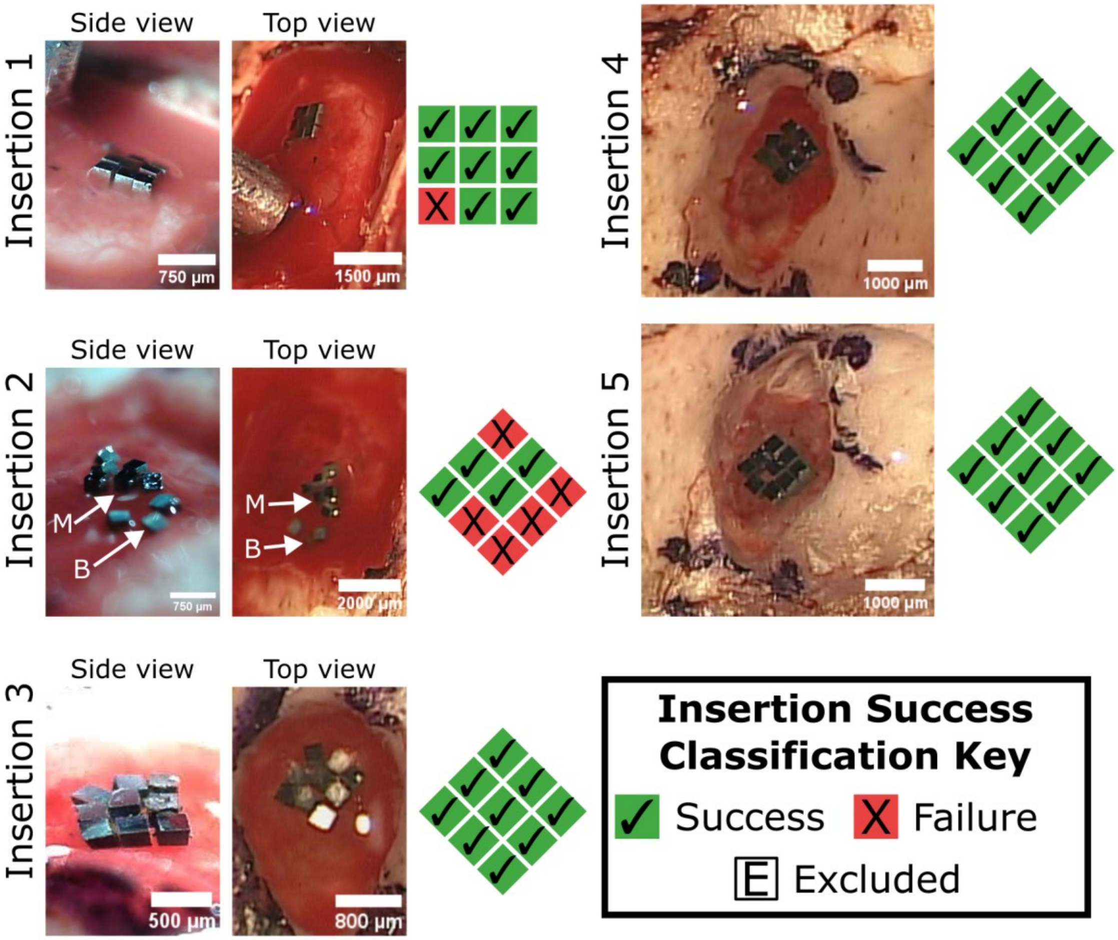


**Figure S4. Outcomes for proof-of-concept mote insertions.** Photos captured after implantation of preliminary mote grids in proof-of-concept insertion experiments. These motes differed slightly from subsequent devices and experiments: these motes were arranged in a 3x3 grid pattern, target fiber length was 500 µm, and the tips were blunt. Additionally, the insertion device was removed in a horizontal motion parallel to the brain surface for insertions 1, 2 and 4, and a mix of horizontal and vertical removal for insertion 3. Out of 45 motes in this dataset, 39 implanted successfully, a success rate of 87%. For each insertion, photos or frames collected post insertion are shown on the left, with a diagram showing insertion success classification on the right.

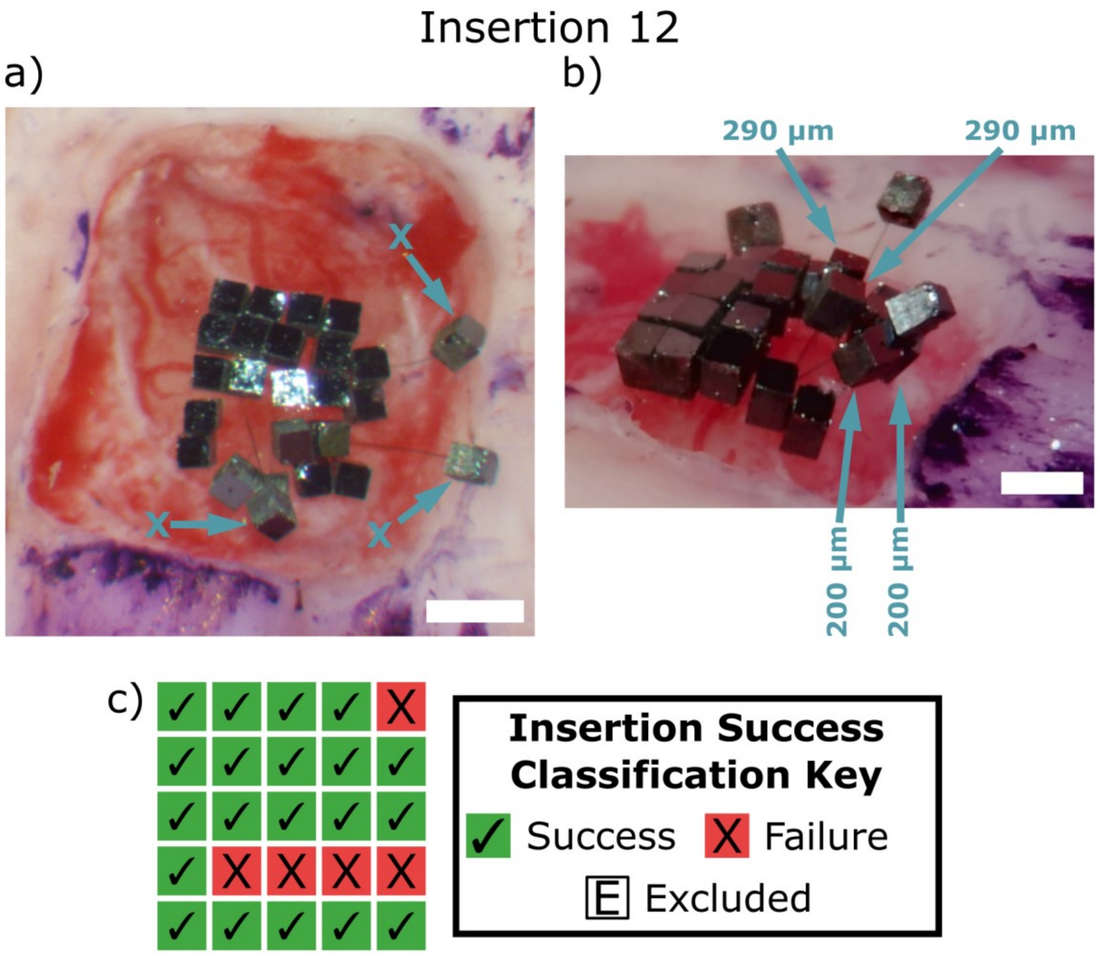


**Figure S5. Illustrative example of an insertion with poor insertional outcomes.** Bird's eye view (a) and left side view (b) of an insertion with poor insertional outcomes shortly after removal of the insertion device. Motes that completely failed to insert (N=3) are labelled with arrows and the letter "X" and are unambiguously identifiable because these motes fell to their side. Motes that mostly inserted are labelled with arrows and the estimated length of fiber still outside of the brain (N=4), rounded to the nearest 10 µm. Two of these motes had more than 200 µm of fiber remaining to be pushed in (N=2) and were therefore classified as failures. c) Diagram showing insertion success classification for this insertion experiment. The layout corresponds to the bird's eye view. (a) Scale bar: 750 µm. (b) Scale bar: 500 µm. Image processing: a) none. b) 10 frames of video captured were averaged, then gamma corrected and contrast adjusted in ImageJ [1].

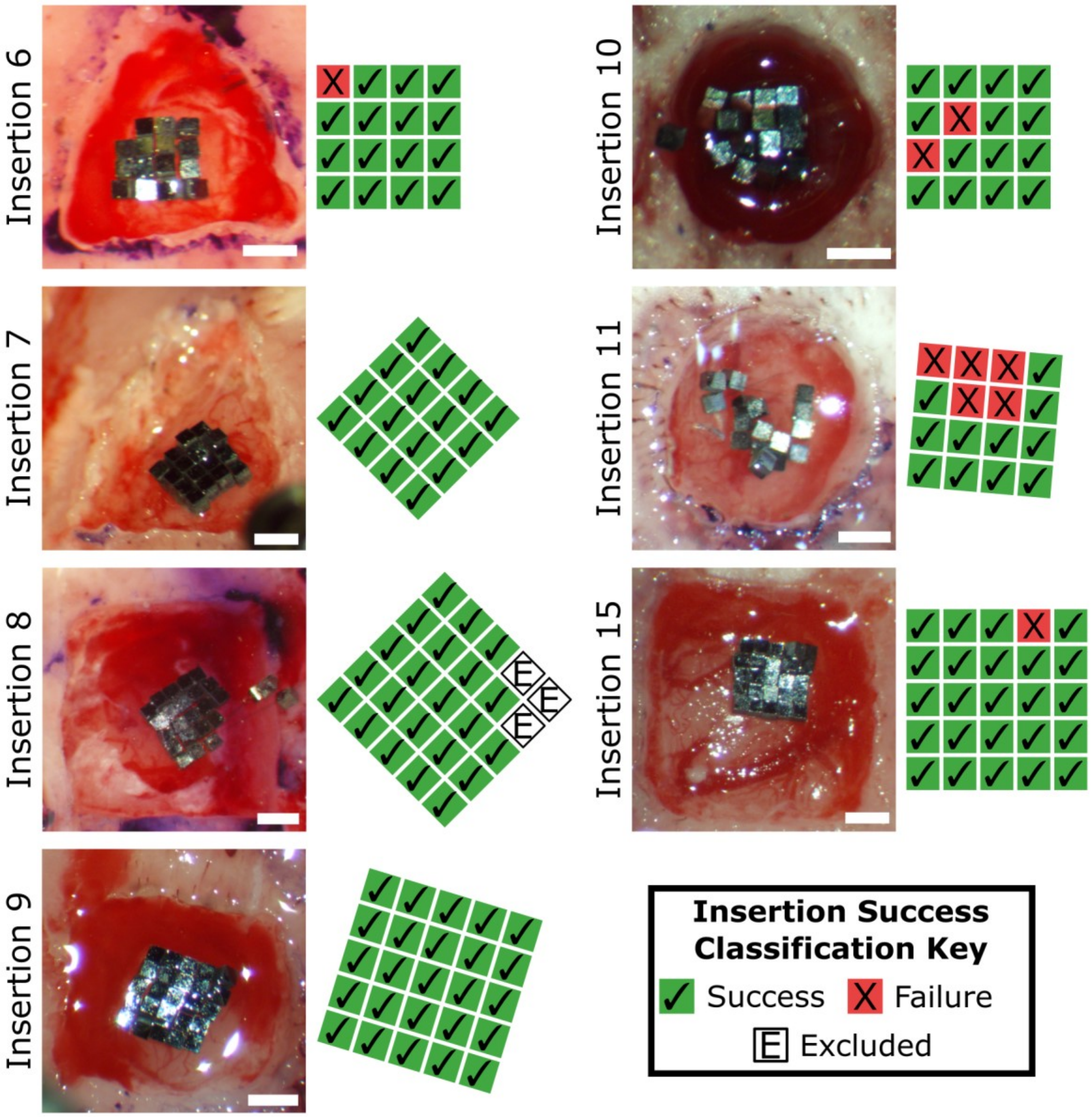


**Figure S6. Outcomes for mote grid insertion experiments.** Photos captured after implantation of 4x4 (N=4) and 5x5 (N=5) mote grids during insertion testing experiments. To reduce redundancy, the outcomes for insertions 14 and 12 are not shown here because they are included in figures 7 and S5, respectively. For each insertion, a bird's eye photo is shown after removal of the insertion device (left) and insertion success classification is shown diagrammatically on the right with layouts matching the bird's eye photos. For insertion 8, three motes were excluded from classification due to surgeon error, as residual dura blocked their insertion into the brain. Scale bars: 750 μm. Note: No image processing was applied to these photos, with the exception of the photo for Insertion 15, which was gamma corrected in ImageJ [1].

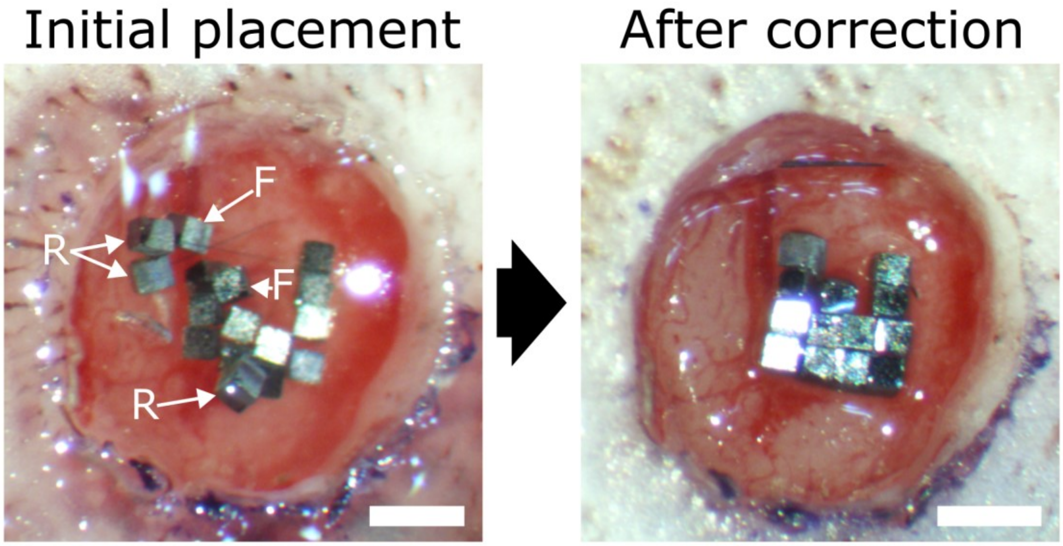


**Figure S7. Example correction of poorly inserted motes.** Bird's eye photos captured of an insertion shortly after the removal of the insertion device (left) and after correction (right). Motes that had completely failed to insert (N=3) were removed via forceps and are labelled "R" (removed) (left). Motes that had mostly inserted with more than 200 µm of non-inserted fiber remaining that were pushed in during correction (N=2) are labelled "F" (fixed) (left). The remaining devices were straightened and, if 0-200 µm of non-inserted fiber remained, pushed in further as part of the procedure. This is from insertion 11, shown in Figure S6. Insertion success classification yielded 11 motes that had successfully inserted (69%), and 5 that failed to insert (19%). Correcting the devices that had mostly inserted increased the proportion of fully inserted motes to 81%. Scale bars: 750 µm.

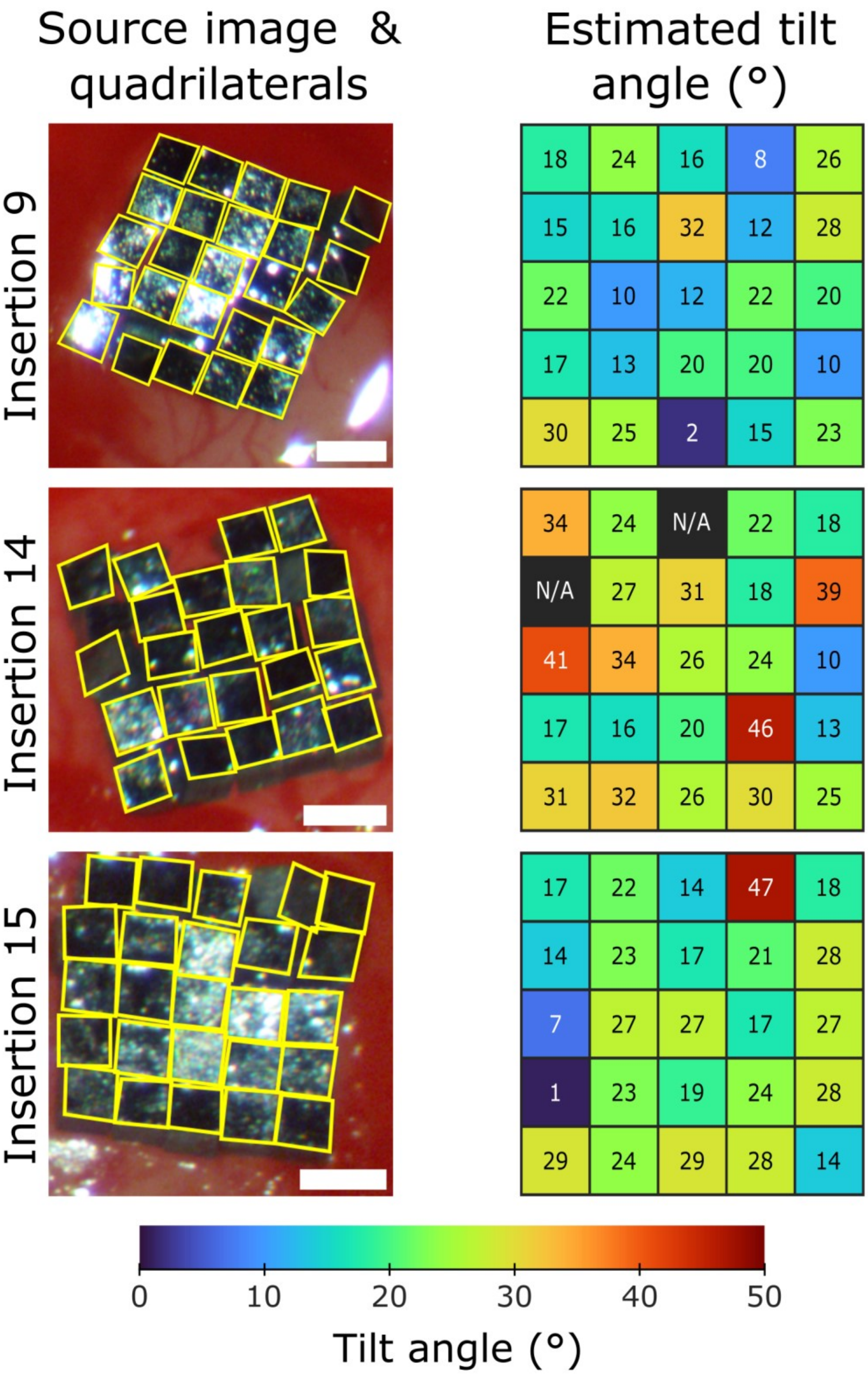


**Figure S8. Visualization of mote tilt angle analysis.** The tilt from horizontal for motes that were implanted onto the surface of the brain was calculated by fitting a quadrilateral to the topside of each mote, using Perspective-n-point calculation to determine the mote pose relative to the camera, and then compared to a reference angle defined by the average pose of all motes implanted in that experiment (N=3 insertions, N=73 motes). On the left, source images are shown with the quadrilaterals (yellow) used for the calculation. On the right, grids with the calculated angles are shown. The layout of motes in the angle grids are the same as in the source images (i.e. the top left mote in the source photo is the same as the top left in the grid, etc.). Scale bars: 400 μm. Image for Insertion 15 was gamma corrected in ImageJ.

| Insertion # | Rat # | Timespan (A/C) | Layout | Reused Motes (#) | Coated in Parylene C (Y/N) | Hemisphere | Craniectomy Coordinates | | Craniectomy Shape | Evaluation | | | Success Rate (%) |
|---|---|---|---|---|---|---|---|---|---|---|---|---|---|
| | | | | | | | M/L | A/P | | S | F | E | |
| 1 | 1 | A | 3x3 | 0 | N | R | 1.0-6.0 | (-8.0)-(-3.0) | R | 8 | 1 | 0 | 89 |
| 2 | 1 | A | 3x3 | 0 | N | L | 0.5-5.0 | (-7.5)-(-2.5) | R | 4 | 5 | 0 | 44 |
| 3 | 2 | A | 3x3 | 0 | N | L | 0.5-3.5 | 1.0-5.0 | O | 9 | 0 | 0 | 100 |
| 4 | 3 | C | 3x3 | 0 | N | R | 0.5-2.5 | 1.5-5.5 | I | 9 | 0 | 0 | 100 |
| 5 | 3 | C | 3x3 | 0 | N | R | 0.5-2.5 | 1.5-5.5 | I | 9 | 0 | 0 | 100 |
| **Total** | - | - | - | - | - | - | - | - | - | 39 | 6 | 0 | 87 |

**Table S1. Proof-of-concept insertion experiment details.** Proof-of-concept insertions with preliminary motes were performed in differing regions of rat cortex with minor differences in technique. Relevant experimental details are listed here for more information on how these experiments differed. Craniectomy coordinates are relative to bregma. Abbreviations: Timespan column: A=Acute, C=Chronic. Hemisphere column: L=Left, R=Right. Craniectomy coordinates column: M/L=Medial/Lateral. A/P=Anterior/Posterior. Craniectomy shape column: R=Rectangle, O=Oval, I=Ice cream cone. Evaluation & success rate columns: S=Success, F=Failure, E=Exception. See Table 1 for details on insertions with larger grid sizes and 1 mm long fibers (Insertions 6-15).

| Failure Mode | # Associations | Primary Category |
|---|---|---|
| Dimpling - blood vessel & swelling | 7 | Brain topography |
| Buckling | 6 | Methodology |
| Early PEG release on much of array | 5 | Methodology |
| Pia hardened by blood | 5 | Surgical complication |
| Brain pushed down by another mote | 4 | Mote interactions |
| Brain blocked by dura | 3 | Surgical complication |
| Evidence of large fiber angle | 3 | Methodology |
| Large fiber angle | 3 | Methodology |
| Mote collision | 3 | Mote interactions |
| Prerelease | 3 | Methodology |
| At edge of swelling | 2 | Brain topography |
| Blood in craniectomy | 2 | Surgical complication |
| Evidence of mote collision | 2 | Mote interactions |
| Horizontal removal of insertion device | 2 | Methodology |
| Blood vessel | 1 | Brain topography |
| Dimpling - blood vessel | 1 | Brain topography |
| Distance to brain | 1 | Surgical complication |
| Evidence of buckling | 1 | Methodology |
| Evidence of loose fiber | 1 | Methodology |
| Evidence of prerelease | 1 | Methodology |
| Evidence of pull out by blood | 1 | Surgical complication |
| Large blood vessel | 1 | Brain topography |
| Loose fiber | 1 | Methodology |

**Table S2. Failure modes for incomplete mote insertions**. Failure modes are listed in descending order from the most associations to the least. Since multiple failure modes were attributed to most motes (19/32), the total number of associations in this table is greater than the number of motes assessed. The third column lists the primary category that was associated with each failure mode.

### Video Legends

**Video 1. Motes readily aggregating in PEG.** Video showing how motes will aggregate in molten PEG. A mote base is placed onto the glass of an insertion device. The soldering iron is then held to the nail shaft and the PEG is melted. Once the mote base is nudged into the molten PEG, it is pulled toward the other mote bases in PEG due to surface tension. The group of motes is prodded with forceps to show how they still stick together when PEG is molten. Bases without fibers were used for demonstration purposes. Video is shown at regular speed. Scale bar: 500 μm.

**Video 2. Mote implantation (vertical view).** Video showing an exemplary mote implantation where all 25 motes in a 5x5 grid successfully implanted into rat cortex. This video was captured from a pen camera looking downward at the craniectomy. The video is sped up by a factor of 8, and much of the saline application step is removed to reduce runtime. Frames from this video are displayed in Figure 6a. Scale bar: 750 μm.

**Video 3. Mote implantation (side view).** Side view of the same implantation shown in Video 2. The video is sped up by a factor of 8, and much of the saline application step is removed to reduce runtime. This video lines up in time with Video 2. Frames from this video are displayed in Figure 6b. Scale bar: 750 μm.

**Video 4. Pushing in Motes That Had Mostly Inserted.** Side view of pushing motes in further with forceps to complete insertion for a 5x5 grid. The device with the large tilt did not push in all the way, but was pushed in deeper. Video is shown at regular speed. Scale bar: 600 μm.
